# Virally encoded single-chain antibody fragments targeting alpha-synuclein protect against motor impairments and neuropathology in a mouse model of synucleinopathy

**DOI:** 10.1101/2024.09.09.612044

**Authors:** Anne-Marie Castonguay, Béatrice Morin, Martin Parent, Thomas M. Durcan, Wen Luo, Irina Shlaifer, Jean-Pierre Julien, Claude Gravel, Martin Lévesque

## Abstract

Parkinson’s disease (PD) is a neurodegenerative disorder mainly characterized by the loss of dopaminergic neurons from the substantia nigra. Affected neurons exhibit intracellular aggregates primarily composed of misfolded and phosphorylated alpha-synuclein (aSyn). In pathological conditions, this presynaptic protein has been shown to be transmitted from cell to cell in a prion-like manner, which contributes to the progression of the disease. Single-chain variable fragments (scFvs) are small polypeptides derived from the binding domains of antibodies that are less immunogenic and have better tissue penetration compared to full antibodies. In this work, we aimed to demonstrate the potential of extracellular scFvs to slow down the propagation of pathological aSyn in an *in vivo* model of synucleinopathy. We generated scFvs that target aSyn, and tested two of them in a PD mouse model consisting of transgenic M83 mice injected with human aSyn pre-formed fibrils (PFFs). The sequence encoding each anti-aSyn scFv was cloned in a self-complementary AAV2 viral vector, and purified particles were administered intravenously. CNS expression of either scFv protected against the development of paralysis and limb weakness, in addition to significantly reducing pathologic aggregates of phosphorylated aSyn in the brain. Moreover, *in vitro* results in human iPSCs-derived dopaminergic neurons suggest that the scFvs can mitigate aSyn spreading by preventing its internalization. Overall, our findings demonstrate that single-chain antibody fragments exhibit strong therapeutic potential in a preclinical mouse model. Thus, our minimally invasive, gene-mediated immunotherapy approach has the potential to serve as an effective treatment for halting the progression of Lewy body diseases.

## Introduction

Parkinson’s disease (PD) is the second most common neurodegenerative disorder after Alzheimer’s disease [1]. It is mainly characterized by the degeneration of dopaminergic neurons from the midbrain [2,3], which is in part responsible for the cardinal motor symptoms of the disease including tremor, bradykinesia, and postural instability [2,4]. Although it was described more than 200 years ago [5], the cause and the exact mechanisms underlying the onset and progression of PD are still unclear. The vast majority of degenerating neurons present intracellular inclusions called Lewy bodies [6,7]. These insoluble aggregates are composed primarily of misfolded and hyperphosphorylated alpha-synuclein (aSyn) [8], a pre-synaptic protein whose functions are still being studied. The discovery that genetic mutations in the gene encoding aSyn (*SNCA*) are directly associated with the onset of the disease in some families confirmed the essential role of this protein in the pathophysiology of PD [9–11].

aSyn is a small intrinsically disordered protein [12]. When it is exposed to various changes in the cellular environment, aSyn can undergo conformational modifications that facilitate aggregation [13]. The results of several experiments in animal and *in vitro* models suggest that aSyn can propagate in the brain and the body as a prion-like protein [14–17]. Indeed, it was shown that aSyn pathology spreads from a site of initiation instead of appearing randomly in various places [18]. The pathology arises either in the periphery or in the brain, and the site of initial pathology determines the course of progression and the main symptoms observed [19–21]. Many now consider PD as a member of the Lewy body diseases family, which also includes dementia with Lewy bodies (DLB), multiple system atrophy (MSA) and pure autonomic failure (PAF). The differential diagnosis is based on the first symptoms observed, which in turn depends on where the synucleinopathy is most abundant [22].

Since Lewy body diseases are proteinopathies, strategies aiming at inhibiting aggregation or limiting cell-to-cell transmission of pathological protein species are widely studied. Immunotherapy is particularly attractive in this regard because it allows the precise targeting of toxic forms of a certain protein. Monoclonal antibodies in the form of immunoglobulin G (IgG) are traditionally the most frequently used immunotherapeutic tools [23]. Several monoclonal antibodies targeting various aSyn strains have been studied [24–30] and some of them are being evaluated in clinical trials [31–37]. However, targeting the brain with IgGs is challenging because of their large size and poor penetration through the blood-brain-barrier [38,39]. To address this challenge, researchers have focused on reducing the size of antibodies while preserving their binding specificity and immune functions. Single-chain variable fragments (scFvs) are composed of the variable regions of the light and the heavy chain of an antibody (V_L_ and V_H_) that are linked together with a flexible and hydrophilic peptide linker. When folded correctly, scFvs can retain the binding specificity of the parental antibody but with a significantly reduced size (∼27 kDa) [40–42]. The small size of the DNA sequences encoding these antibodies allows for their incorporation into an expression cassette that fits within an adeno-associated viral (AAV) vector, an attractive tool among gene therapy applications [43]. Another notable advantage of scFvs over full antibodies is their reduced immunogenicity due to their smaller size, and absence of the fragment crystallizable region (Fc), which interacts with immune cells and molecules [43]. In a therapeutic setting, they can be used to modulate (e.g., blocking) the function of their target(s), act as agonists or antagonists for receptors, inhibit aggregation and cell entry, or stimulate degradation of their target(s) [44,45].

In this work, we report two scFvs derived from novel antibodies targeting the C-terminal part of aSyn that show great potential in reducing pathological aSyn propagation in a synucleinopathy mouse model. Intravenous administration of AAV vectors encoding the secreted scFvs was performed in transgenic mice overexpressing human A53T aSyn (M83 line) that were previously inoculated in the dorsal striatum with human aSyn pre-formed fibrils (PFFs). The production of either scFvs against aSyn in the CNS significantly protected against motor impairments and hind limb paralysis compared to our control scFv. Accordingly, we observed reduced pS129-aSyn pathology in the brain of the mice that received scAAVs encoding the scFvs against aSyn. Contrary to most passive immunotherapy approaches, our strategy requires a single non-invasive injection for efficient brain delivery. Hence, we believe our approach could be adapted for human trials and hopefully slow disease progression to maintain the quality of life of people living with Lewy body diseases such as Parkinson’s.

## Results

### Monoclonal antibody generation and characterization

We first generated novel monoclonal antibodies by immunizing mice with a phosphorylated (S129) peptide corresponding to a portion of the C-terminal region of human aSyn (**Figure 1A**). We chose to target the carboxy-terminal region because it contains the serine 129 that is found phosphorylated in pathological aggregates, and because studies have shown this region to still be accessible to antibodies when the protein is aggregated [46]. Splenocytes were harvested from three mice that developed the strongest positive immune response against the peptide and were fused to murine NS0 myeloma cells to generate hybridomas [47]. Hybridoma clones that positively bound to the phosphorylated peptide in ELISA (∼20 clones) were further tested to ensure specificity. Hybridoma supernatants were used as primary antibody solutions in western blot experiments on HEK cell lysates previously transfected with an aSyn expression plasmid in combination or not, with the polo-like kinase 2 (PLK2) expression plasmid. PLK2 has been shown to phosphorylate aSyn on the serine 129 *in vivo* [48]. Cells transfected with an expression plasmid encoding the human mutant S129G-aSyn protein, which cannot be phosphorylated by PLK2, were also used. Non-transfected cells served as negative control. We selected 2 IgG antibodies (73B3 and 73D1) that showed specificity to pS129-human aSyn and 2 IgGs (826B2 and 833D4) that could bind to both pS129- and non-phosphorylated human aSyn (**Figure 1B**). To ensure that the antibodies could bind non-denatured native human aSyn, we also used hybridoma supernatants as a primary antibody solution in immunofluorescence on our transfected HEK cells using the same conditions as previously described. All 4 antibodies demonstrated the same specificity in immunofluorescence as was observed in western blots (**Figure 1C**).

**Figure 1:**
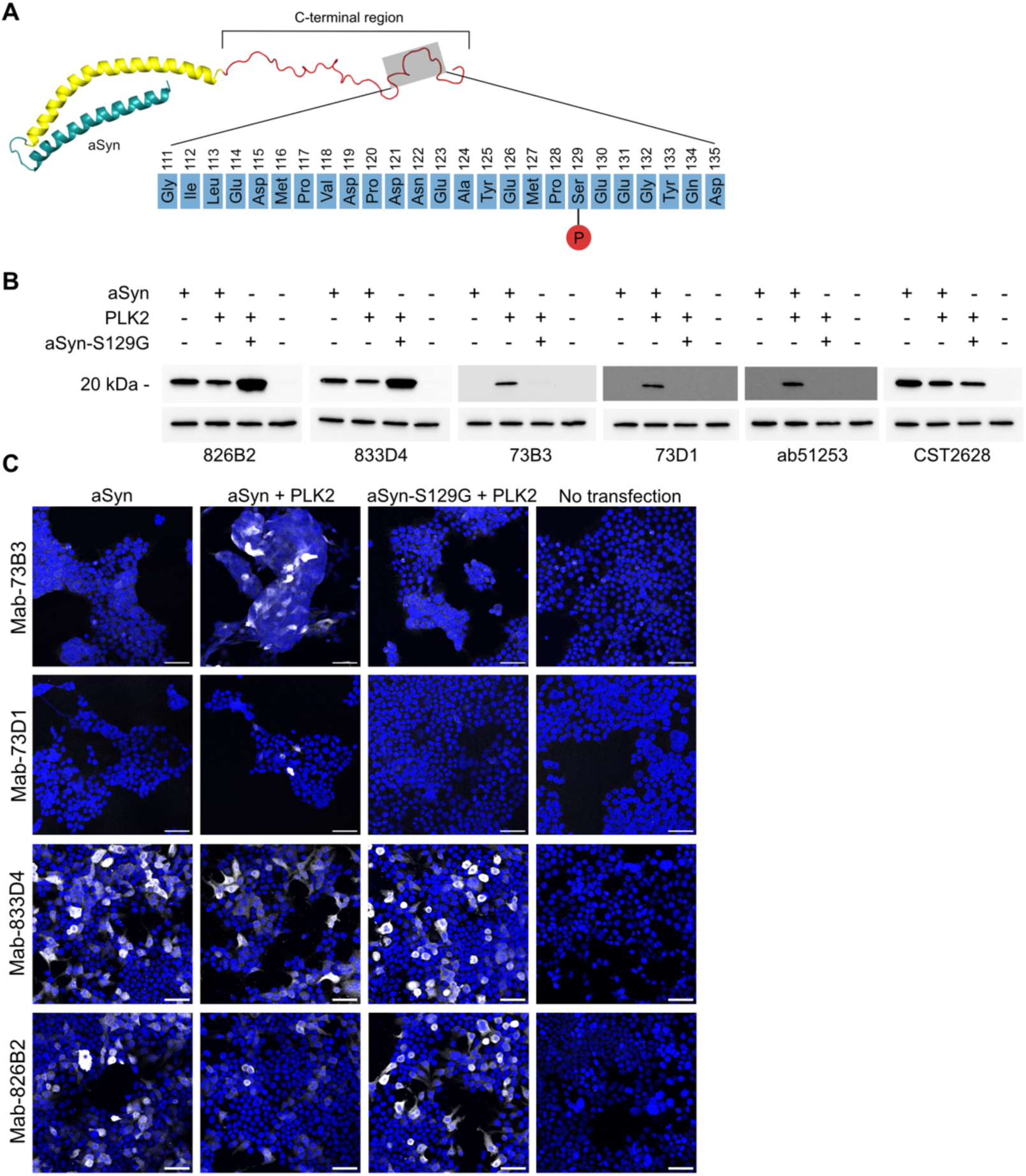
Generation of murine monoclonal antibodies against total and phosphorylated human aSyn. A) The 25 a.a. peptide used to immunize the mice and location in the full-length protein. B) Western blot showing the specificity of each hybridoma clone selected on HEK cells lysate. Cells were transfected with either an aSyn encoding plasmid alone or in combination with a PLK2 kinase encoding plasmid. A plasmid encoding the mutated aSyn-S129G that cannot be phosphorylated at the serine 129 was also used. A condition with no transfection was used as a negative control. Commercial antibodies against total and pS129-aSyn were used as comparison (ab51253 against pS129-aSyn, and CST2628 against total aSyn). C) Representative images of immunofluorescence on HEK cells that were transfected with the plasmids described above using each hybridoma clone media as a primary antibody solution. Antibody-specific signal is shown in white, DAPI is shown in blue. Scale bar = 50μm.

The four selected IgGs were purified from hybridoma culture media using Protein G agarose beads. They were tested on fixed brain sections from patients diagnosed with PD, MSA, and DLB to detect pathological aSyn inclusions (**Supplementary table 1**). Age-matched control brain sections were also used. Both phospho-specific antibodies (73B3 and 73D1) recognized aSyn inclusions in sections from the substantia nigra (SN) of diagnosed PD patients, but not in SN sections from control subjects (**Figure 2A and Supplementary table 2**). Detected structures include perikaryal inclusions, Lewy body-like aggregates and dystrophic neurites. Both pan-aSyn-specific antibodies (833D4 and 826B2) also recognized aSyn inclusions in the SN of PD patients, although they also showed some basal staining in control brains. This is consistent with the fact that those antibodies also recognize non-phosphorylated aSyn. The 73B3 and 73D1 antibodies also bound aSyn inclusions in the cortex of MSA and DLB patients, and the staining pattern resembled the one from the commercial antibody used as a control for anti-pS129-aSyn staining (BioLegend #825701) (**Figure 2B and Supplementary table 2**). The antibodies 833D4 and 826B2 recognized inclusions of aSyn in the cortex of DLB-, but not MSA-diagnosed patients. The 833D4 and the 826B2 overall stained fewer aggregates compared to the pS129-aSyn-specific antibodies in similar brain sections from the same patients (**Figure 2B and Supplementary table 1**). These results indicate that our novel monoclonal antibodies can detect pathological aSyn inclusions in the brains of patients suffering from Lewy body diseases.

**Figure 2:**
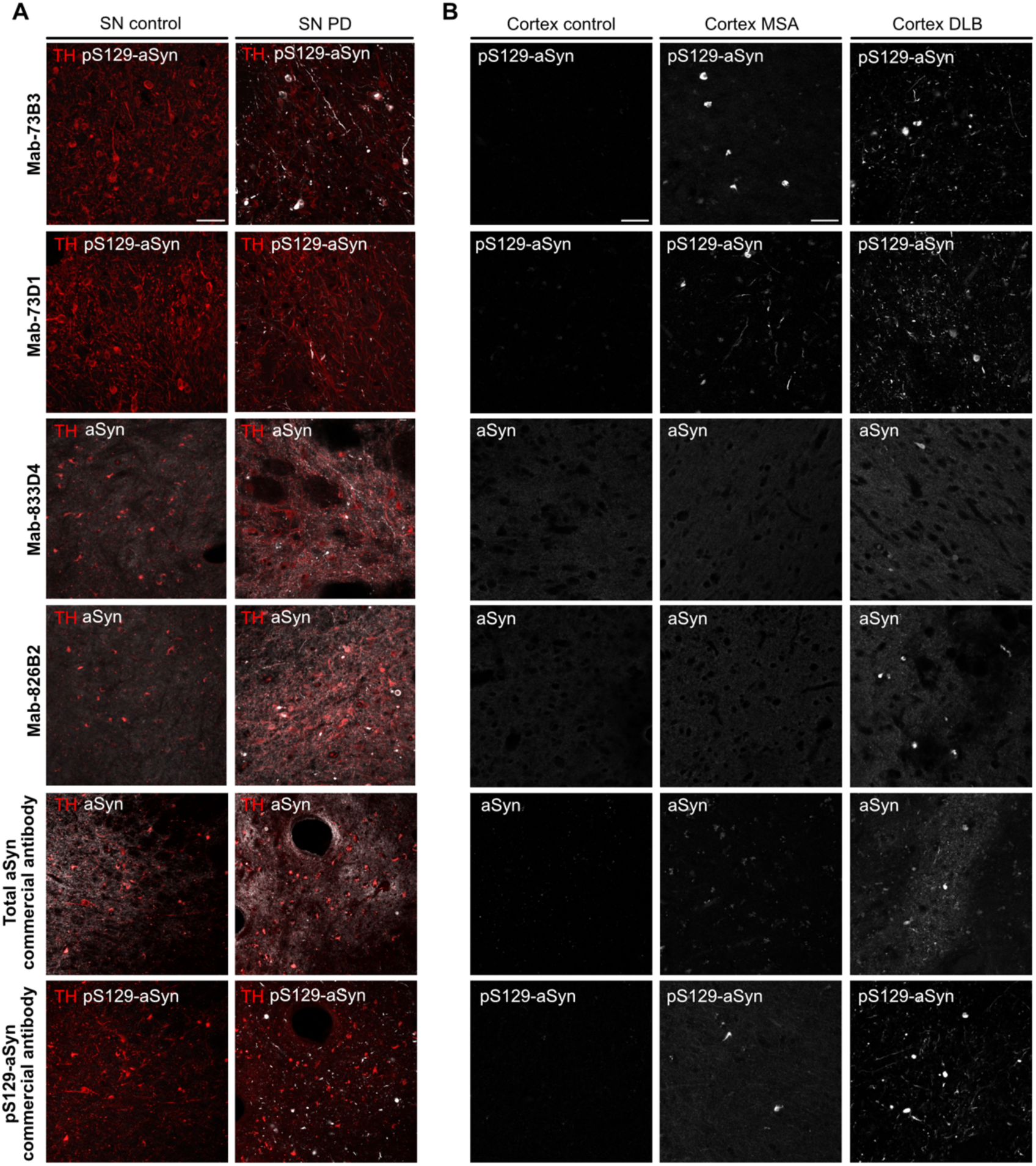
Novel monoclonal antibodies recognize pathological aSyn inclusions in brain sections from patients suffering from PD, DLB and MSA. A) Representative images of immunofluorescence using our purified monoclonal antibodies on human Parkinson’s disease (n=3), and control (n=2) substantia nigra sections. Antibody-specific signal is shown in white while TH is shown in red. Commercial antibodies were used as a comparison (CST2628 against total aSyn, and BioLegend 825701 against pS129-aSyn). B) Representative images of immunofluorescence using our purified monoclonal antibodies on human MSA (n=3), DLB (n=3) and control (n=2) cortex sections. SN = Substantia nigra; MSA = multiple system atrophy; DLB = dementia with Lewy bodies. Scale bar = 100μm.

### Monoclonal antibodies 833D4 and 73B3 demonstrate high affinity to aSyn and pS129-aSyn and can mitigate their aggregation *in vitro*

We focused on these two antibodies to compare the therapeutic potential of a pS129-aSyn-specific antibody (73B3) versus a pan-aSyn-specific antibody (833D4). Using surface plasmon resonance (SPR), we characterized the affinity and binding kinetics of each antibody towards human aSyn and aSyn phosphorylated at S129 (pS129-aSyn). Based on equilibrium dissociation constants (KD), both antibodies displayed high affinity (in the nanomolar range) for their target [49]. The 833D4 showed slightly higher affinity for pS129-aSyn compared aSyn (KD^aSyn^= 2.60nM, KD^pS129^= 0.91nM) with very slow dissociation rates for both targets (**Figure 3 A, B and E**). Similarly, 73B3 exhibited strong affinity for pS129-aSyn (KD^pS129^= 1.66nM) but bound to aSyn with a five-fold lower affinity (KD^aSyn^= 8.25nM) (**Figure 3 C-E**). Importantly, the response (resonance units, proportional to the number of bound analytes) of 73B3 towards aSyn was lower, with a much faster dissociation rate compared to the phosphorylated protein, highlighting its stronger specificity for pS129-aSyn (**Figure 3 C-E**).

**Figure 3:**
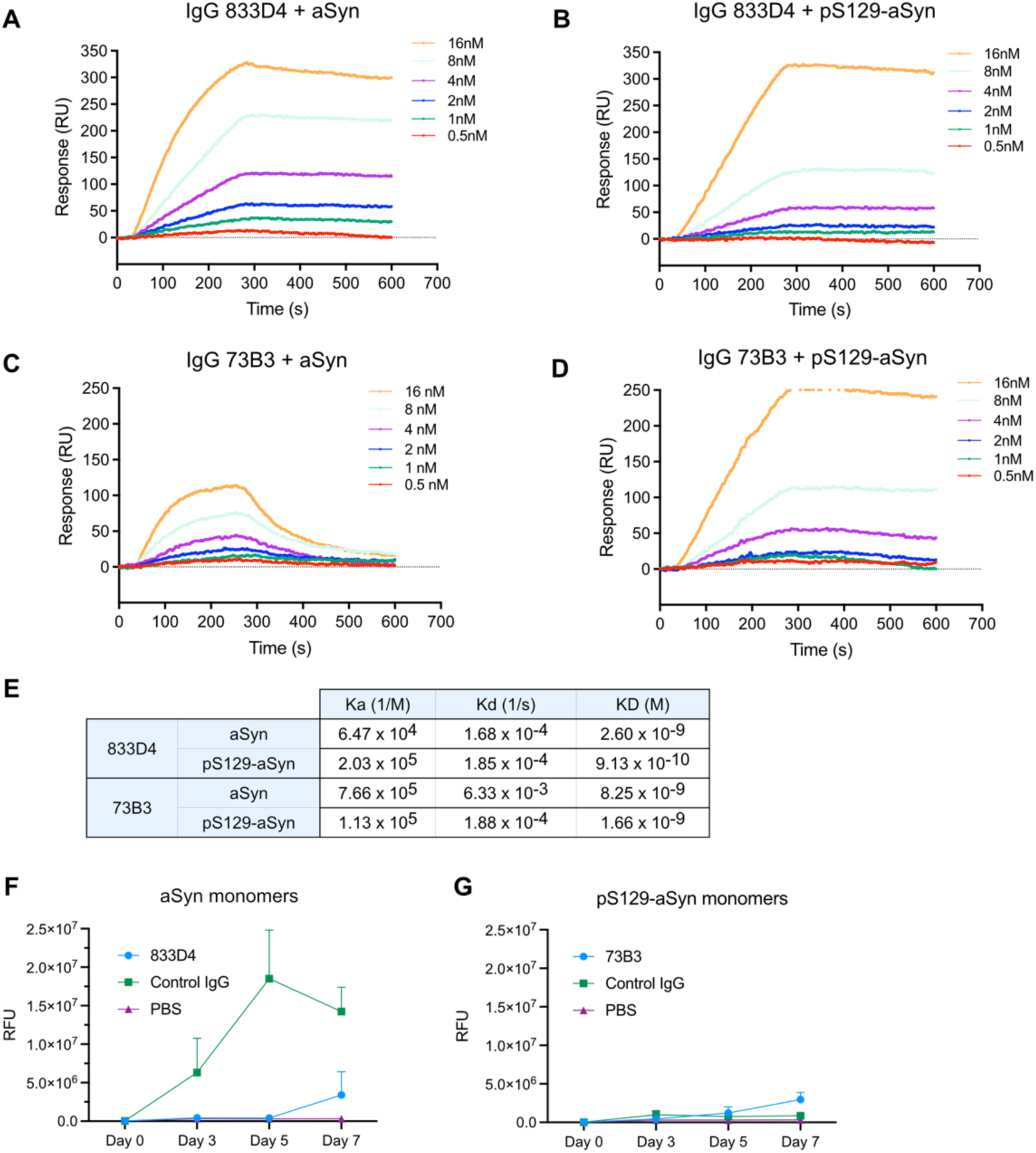
IgG 833D4 and 73B3 have a high affinity to aSyn and pS129-aSyn monomers and mitigate fibrillization. A-D) Real-time binding curves of 833D4 and 73B3 IgGs to aSyn and pS129-aSyn measured using surface plasmon resonance (SPR). aSyn monomers were coated on the sensor chip, and the antibodies were flown over at different concentrations. The response is shown in resonance units, which is proportional to the number of bound antibodies to the chip. E) Association (Ka), dissociation (Kd) and equilibrium (KD) constants of both antibodies towards aSyn and pS129-aSyn. F) and G) A Thioflavin T (ThT) aggregation assay was performed by incubating aSyn monomers in the presence of ThT with agitation for seven days at 37°C (n=3). F) Relative fluorescence units (RFU) of ThT correlating to the level of β-sheets structures of aSyn monomers in the presence of 833D4 or an unspecific IgG. G) RFU of ThT with pS129-aSyn monomers in the presence of 73B3 or an unspecific IgG.

To verify if our antibodies could prevent *in vitro* fibrillization of aSyn, we performed a Thioflavin T aggregation assay using aSyn or pS129-aSyn monomers in the presence of IgG 833D4, 73B3, or an unspecific antibody (IgG2a against La Crosse virus G1 glycoprotein). The 833D4 antibody prevented fibrillization of aSyn monomers during the 7 days of the test, while the non-specific IgG did not (**Figure 3F**). pS129-aSyn monomers did not form fibrils as efficiently as aSyn monomers in this assay; a fact that was previously reported [50]. We thus could not evaluate the anti-aggregation properties of the 73B3 IgG in this assay (**Figure 3G**). These results indicate that the IgG 833D4 can inhibit the *in vitro* fibrillization of aSyn monomers.

### Single-chain variable fragments (scFvs) generation and validation

ScFvs are composed of the variable regions of the light (Vk) and the heavy (VH) chain of a monoclonal antibody (**Figure 4A**). To synthesize them, the entire sequences of the light and heavy chains of our antibodies were obtained by next-generation sequencing and scFv-encoding synthetic cDNAs were assembled in the following order: IgG heavy chain-encoded variable region with its secretion signal, (Gly_4_Ser)_4_ flexible linker, IgG light chain-encoded variable region, and a small Myc tag encoding region. These synthetic scFv-encoding genes were then cloned in a self-complementary AAV (scAAV) cloning vector plasmid (pscFv) with a CMV promoter/enhancer. A sequence encoding a combined WPRE-SV40 polyadenylation signal (WpPA) [51] was present downstream of the scFv-encoding open reading frame for increased expression. Importantly, the signal peptide from the parental IgG heavy chain was kept, enabling secretion in the extracellular space. These scFv-encoding plasmids were first used to transfect HEK cells for scFv production and analysis, and were subsequently encapsidated into scAAV2/B10- or scAAV2/retro-bearing virions for *in vivo* and *in vitro* use, respectively (**Figure 4B**). A scAAV2 encoding a scFv against GFP [52] (termed control scFv here) was used as a negative control in our *in vitro* and *in vivo* experiments, as no GFP was expressed in any of our models.

**Figure 4:**
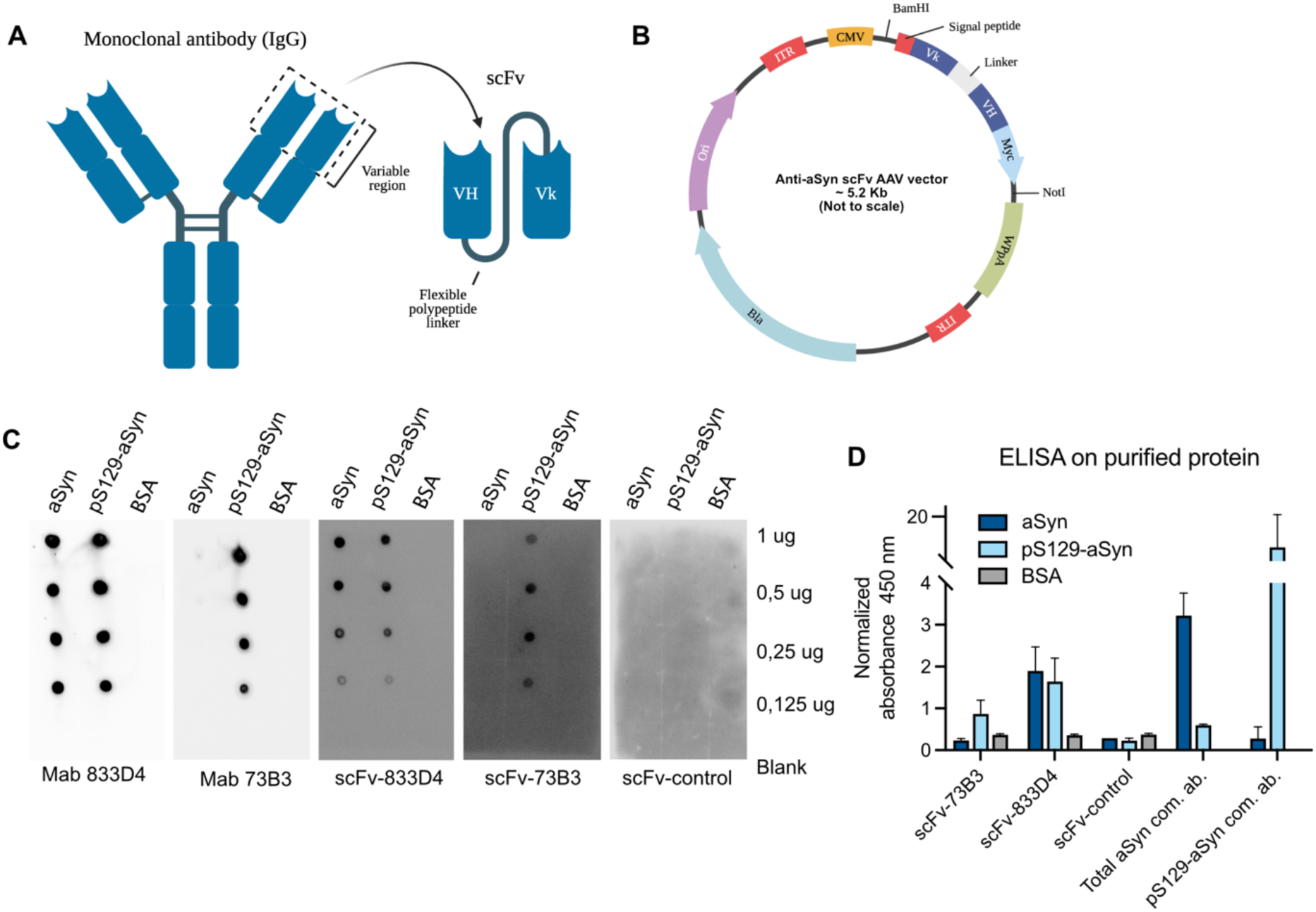
The scFvs generated from our monoclonal antibodies retained their binding specificity. A) Illustration of the structure of a scFv generated from a monoclonal IgG. B) Illustration of the scFv construct cloned inside a scAAV expression plasmid. C) Dot blot performed by blotting aSyn or pS129 aSyn monomers (and BSA as a negative control) on a nitrocellulose membrane. Membranes were either incubated with cell media from HEK cells previously transfected with each pscFvs, or with the purified monoclonal IgGs. Results show that both scFvs maintained the same specificity as their parental IgG. D) ELISA performed by coating the plate with purified monomers of aSyn or pS129-aSyn (or BSA as a negative control) showing the specificity of scFv-833D4 and scFv-73B3. Again, the cell media from HEK cells previously transfected with each pscFvs was used, and commercial antibodies to total (CST2628) and pS129-aSyn (Wako 015-25191) were used as positive controls (n=2).

To verify whether the scFvs retained the binding specificity of the IgGs they were derived from, we transfected HEK cells with the plasmids encoding the scFvs and used the culture media from the transfected cells as primary antibody solution in dot blot and ELISA assays on purified aSyn and pS129-aSyn protein. Bovine serum albumin (BSA) was used as a negative protein control for both experiments, and commercial antibodies to pan and pS129-aSyn were used as positive controls. In the dot blot assay, the scFv-833D4 and scFv-73B3 showed specific binding to purified aSyn and pS129-aSyn, respectively (**Figure 4C**). We obtained comparable results in the ELISA on purified aSyn protein, with strong positive binding on total aSyn for the scFv-833D4, and pS129-aSyn for the scFv-73B3 (**Figure 4D**). Given that these experiments were conducted using scFv-containing cell supernatant, the signal intensity cannot be directly correlated with binding affinity, as the precise concentration of scFv in the medium was undetermined. These results confirmed that the scFvs 833D4 and 73B3 maintained the binding specificity of their parental IgG.

### The injection of scAAVs encoding secreted anti-aSyn scFvs significantly protects against motor impairments and aSyn neuropathology in a synucleinopathy mouse model

To evaluate the therapeutic potential of our scFvs, we used hemizygous M83 transgenic mice aged between 60 and 90 days. This mouse line overexpresses human A53T-aSyn under the control of the murine prion protein (PrP) promoter. M83 mice develop age-dependent motor symptoms leading to paralysis and death, accompanied by intracytoplasmic neuronal aSyn inclusions mimicking aggregates found in the brain of PD patients [53]. To accelerate the pathological aSyn aggregation and spread, we injected human aSyn PFFs in the right dorsal striatum as described previously [54]. As mentioned previously, the anti-aSyn scFvs were encapsidated using a Cap-B10-encoding helper plasmid. This recombinant AAV capsid confers the vector the capacity to pass the blood-brain barrier and to infect neurons in mice and marmosets [55]. Using this serotype thus allows minimally invasive administration and increases translatability to the clinic.

Intravenous injection of scAAVs encoding secreted scFv-833D4, scFv-73B3 or the control scFv was performed into the retro-orbital sinus 7 days after intrastriatal injection of PFFs (or PBS as a negative control). Starting 8 weeks post-PFFs injection, mice were closely monitored with a scoring system (detailed in **Supplementary material)** to assess the onset of motor symptoms (**Figure 5A)**. Mice that received intrastriatal PFFs and the scAAV-scFv-control developed progressive motor impairments, beginning with limb weakness and gait defects (between 9 and 12 weeks post-PFFs), and generally progressed to complete paralysis within a week of symptoms onset. This progression was associated with a significantly higher clinical score at the time of sacrifice compared to the mice injected with scAAVs encoding scFv-833D4 or scFv-73B3 (**Figure 5B**). Notably, scFv-833D4 and scFv-73B3 treatments delayed the onset of symptoms compared to the control scFv (**Figure 5C**). We performed rotarod and a grip strength test before sacrificing the mice at a maximum of 12 weeks post-PFFs injection. On the rotarod assay, mice injected with PFFs and the scAAV-scFv-833D4 had a significantly longer latency to fall compared to mice injected with the scAAV-scFv-control, who showed impaired coordination. There is a strong indication that injection of scAAV-scFv-73B3 also ameliorates the rotarod performance, but it did not reach significance because of high intra-group variability (**Figure 5D**). In the grip strength test, both scAAV-scFv-833D4 and scAAV-scFv-73B3 significantly protected against the loss of strength compared to scAAV-scFv control (**Figure 5E**).

**Figure 5:**
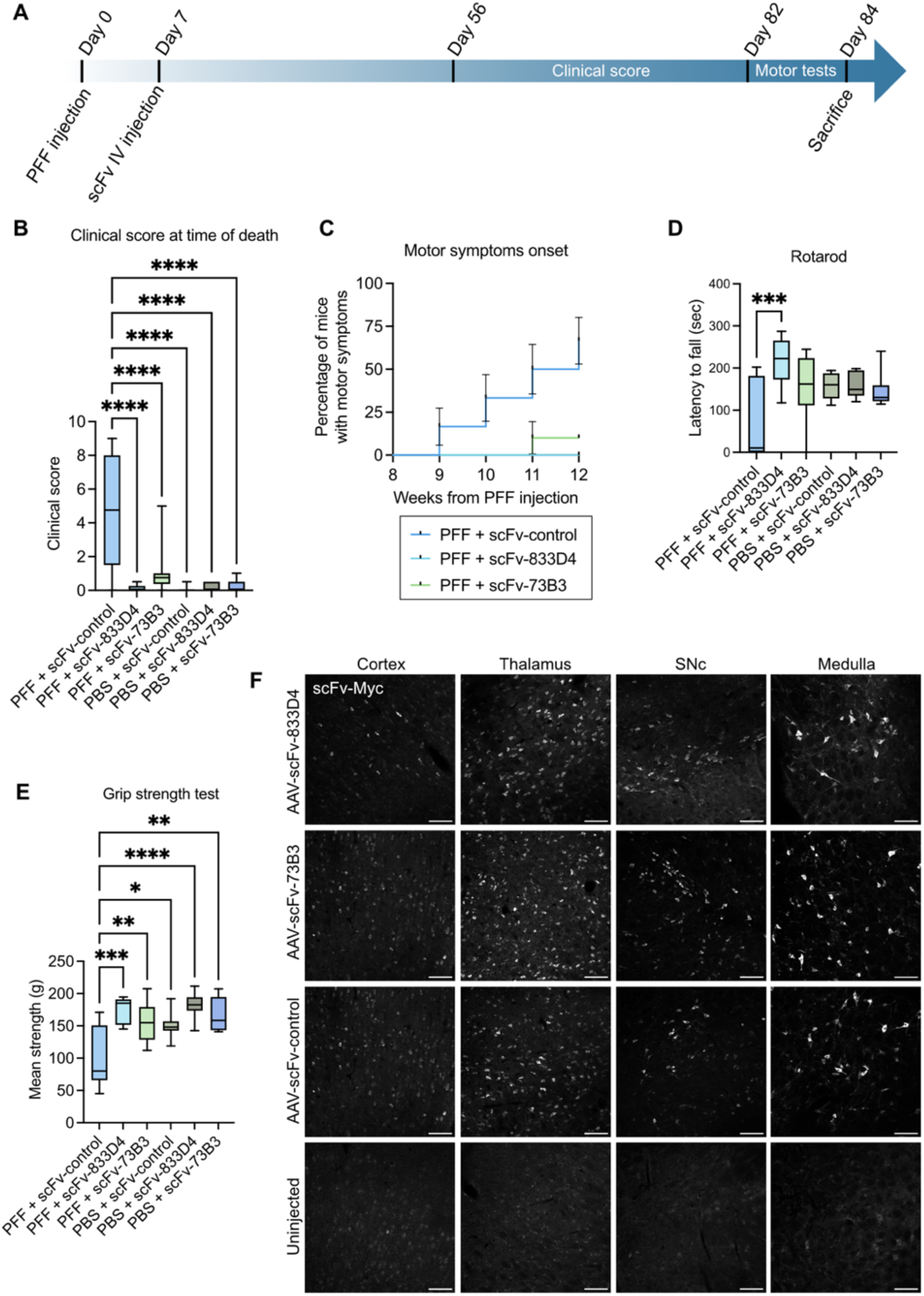
Intravenous injection of scAAV2/B10 encoding scFv-833D4 and scFv-73B3 allows brain expression and protects against motor impairments in the M83 with intrastriatal PFFs mouse model. A) Experimental timeline. B) Clinical score at time of sacrifice in mice that received intrastriatal PFF or PBS with subsequent IV injection of scAAV-scFv. C) Percentage of mice in each PFF-injected group showing motor symptoms (clinical score > 3) as a function of the number of weeks from PFFs injection. D) Latency to fall on the rotarod test. E) Mean strength obtained in the grip strength test. For B to E, n=10-12/group, One-way ANOVA with Dunnett’s multiple comparison test *p<0.05, **p<0.01, ***p<0.001, ***p<0.0001. F) Representative images of a Myc tag immunofluorescence showing brain expression of the scFv-833D4, scFv-73B3 and scFv-control 12 weeks after intravenous injection, compared to a non-injected mouse. Scale bar = 100 μm.

Twelve weeks following PFFs injection (or at the time when mice reached a clinical score of 9), mice were sacrificed, and histological analysis was performed. We first ensured that the anti-aSyn scFvs or control scFv were expressed in the brain of the injected mice with immunofluorescence against the human Myc tag present at the C-terminal of each scFv. Successful brain expression was observed in various regions, with the highest signal observed in neurons of the thalamus and medulla (**Figure 5F**). This result confirmed that the recombinant scAAV2/B10 injected intravenously effectively infected brain cells in our model. No specific anti-Myc epitope immunofluorescence signal was observed in brain cells of PFF-injected M83 mice that did not receive a scAAV injection. Consistent with the AAVCap-B10 published properties [55], low amounts of anti-Myc tag immunofluorescence were observed in the liver (**Supplementary fig. 1**).

At the time of sacrifice in mice that received the scAAV-scFv-control one week after intrastriatal PFFs, numerous inclusions of pS129-aSyn were found in various brain regions, particularly in the midbrain, pons, and medulla (**Figure 6A**). Lewy body-like inclusions were also abundant in the spinal cord, which could explain the severity of the motor symptoms in this group (**Supplementary fig. 2A**). These aggregates exhibited positive staining for β-sheet structures, as illustrated with β-sheet staining (Amytracker© dye, **Supplementary fig. 2B**), indicating a Lewy body-like phenotype. In contrast, M83 mice that received intrastriatal PBS instead of PFFs do not display any brain pathology at sacrifice (**Supplementary fig. 3**), which is to be expected at this young age. Injection of either scAAV-scFv-833D4 or scAAV-scFv-73B3 significantly reduced pS129-aSyn pathology in the brain of M83 mice that received intrastriatal PFFs compared to the scAAV-scFv-control, indicating protection against the pathological spread of aSyn (**Figure 6B-E**). This result was confirmed in western blots from brain lysates, where we observed significantly less pS129-aSyn relative to total aSyn in the medulla of the mice that received either anti-aSyn scFvs (**Figure 6F-G**). Moreover, the total aSyn protein level was not different between groups (**Figure 6F,H**). Histological analysis revealed no significant loss of tyrosine hydroxylase-positive (TH^+^) neurons in the SNc or TH^+^ axons in the striatum of any experimental group (**Supplementary fig. 4 A-F**). To ensure that the control scFv did not have any biological effect in our model, we injected a control group with saline instead of the control scFv. No major differences were observed between these two groups in terms of motor phenotype or brain pathology, confirming that the control scFv does not influence the pathological process in our model (**Supplementary fig. 5 A-M**). These results suggest that the virally encoded anti-aSyn scFvs can significantly decrease aSyn brain pathology and motor dysfunctions in an aggressive synucleinopathy mouse model.

**Figure 6:**
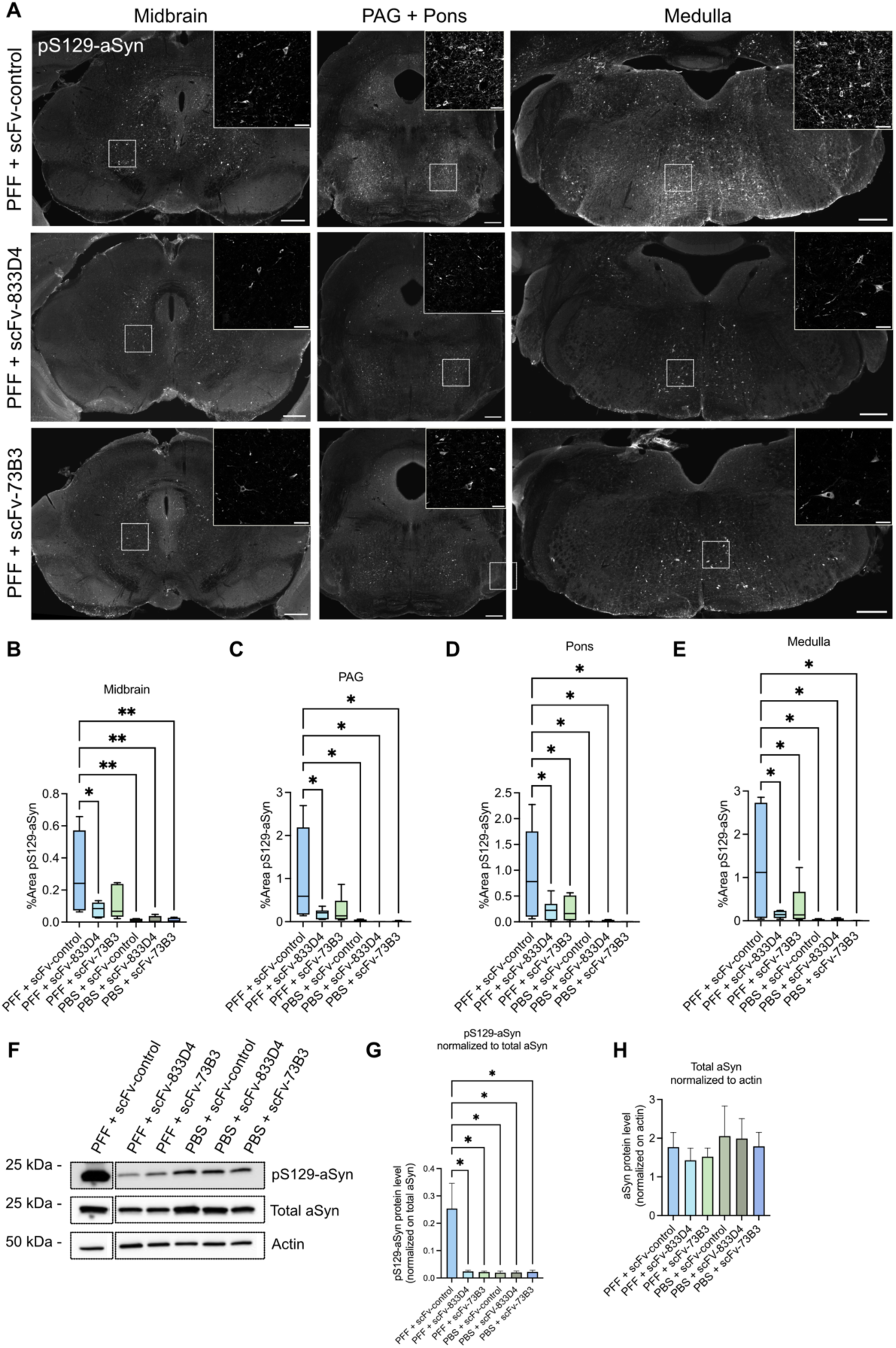
Intravenous injection of scAAVs encoding scFv-833D4 and scFv-73B3 in M83 mice reduce the burden of pS129-aSyn inclusions 12 weeks after PFFs intrastriatal injection. A) Representative images of pS129-aSyn immunostaining in the midbrain, PAG/pons and medulla of M83 mice injected with PFFs and with scAAVs encoding scFv-833D4, scFv-73B3, and scFv-control. B), C), D) and E) Quantification of the pS129-aSyn staining as a percentage of the area measured (n=6/group). One-way ANOVA with Dunnett’s multiple comparison test *p<0.05. **p<0.01. F) Representative images of pS129-aSyn and total aSyn in western blot on medulla tissue lysate. G) Quantification of pS129-aSyn normalized on total aSyn. One-way ANOVA with Dunnett’s multiple comparison test *p<0.05. H) Quantification of total aSyn normalized on actin (n=3-4/group). One-way ANOVA with Dunnett’s multiple comparison test. (n=3-4/group).

### scFv-833D4 and scFv-73B3 are internalized by neurons and microglia *in vivo*, and interfere with entry of exogenous aSyn preformed fibrils in human iPSCs-derived dopaminergic neurons in vitro

To investigate if the scFvs can be internalized by non-producing cells, we used an AAV vector construct (AAV-scFv-IRES-GFP) encoding both our secreted HA-tagged scFv 833D4, and a non-secreted GFP (**Figure 7A**). This was made possible with the use of two open-reading frames (ORFs) controlled by the same CMV promoter by inserting a sequence encoding an internal ribosomal entry site (IRES) between the two ORFs. This vector was encapsidated with the AAVCap-B10 capsid, purified and injected intravenously into aged homozygous M83. Three weeks after the IV injection, the mice were sacrificed, and the distribution of scFv-833D4, pS129-aSyn, and GFP was assessed on fixed brain sections. While most scFv-immunoreactive cells were found to be also GFP-positive, some cells were found scFv-positive but GFP-negative, suggesting they were not infected by the viral vector (**Figure 7B**). Moreover, we also observed colocalization of the scFv and pS129-aSyn in some GFP-negative neurons (**Figure 7C**). Because the morphology of the observed scFv-positive cells was varied, we evaluated if the scFv could also be present in microglia by performing Iba1 immunostaining. We indeed observed frequent colocalization of the scFv in Iba1-positive cells (**Figure 7D**) in the absence of GFP fluorescence. These results suggest that the secreted scFvs can be internalized by neurons and microglia, which could contribute to aSyn clearing.

**Figure 7:**
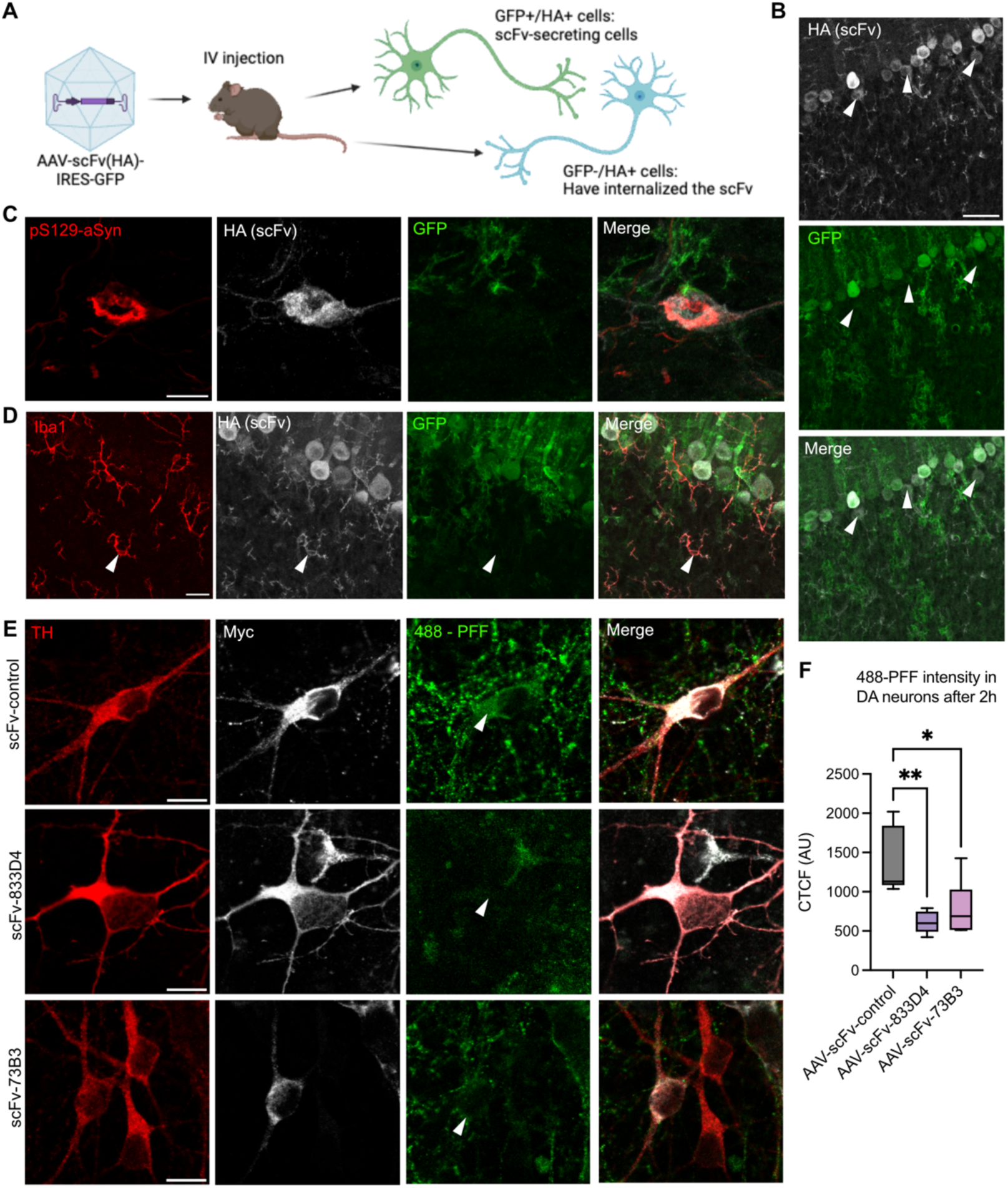
Secreted anti-aSyn-scFvs are internalized by non-secreting cells and prevent PFFs internalization in human iPSCs-derived dopaminergic neurons. A-C) Aged M83 homozygous mice were injected intravenously with an AAV-CapB10-scFv-IRES-GFP. This construct allows the expression of a secreted HA-tagged anti-aSyn scFv (833D4) and a non-secreted GFP protein. Mice were sacrificed 3 weeks post-injection. B) Representative confocal image taken in the cerebellum showing HA (scFv) in white and GFP in green. Red arrows point to neurons that are HA^+^/GFP^-^ and thus have internalized the scFv. C) Confocal image showing the colocalization of the scFv (HA) and pS129-aSyn in a GFP-negative neuron from the medulla. D) Confocal images showing the colocalization of the scFv (HA, white) and Iba1 (red), indicating the internalization of scFvs by microglia. E) Representative images of human iPSCs-derived dopaminergic neurons (identified with a TH staining) infected with a scAAV2/retro encoding either scFv-833D4, scFv-73B3 or control scFv (visualized with Myc tag staining) and exposed for 2 hours to human aSyn PFFs tagged with Alexa Fluor 488. Scale bar = 10μm. F) Quantification with corrected total cell fluorescence (CTCF) of Alexa Fluor 488 intensity inside TH+ neurons (n=6 coverslips/condition, from 3 independent differentiations). One-way ANOVA with Dunnett’s multiple comparison test, *p<0.05, **p<0.01.

To evaluate whether scFv-833D4 and scFv-73B3 can modulate aSyn spreading by inhibiting the internalization of extracellular aSyn, we utilized human aSyn preformed fibrils (PFFs) tagged with Alexa Fluor 488™ (488-hPFF). Human induced pluripotent stem cell (hiPSC)-derived dopaminergic neurons with a triplication of the SNCA gene (3xSNCA) [56] were used, as they consistently develop aggregates of pS129-aSyn following exposure to exogenous human PFFs (data not shown). Mature hiPSC-derived dopaminergic neurons (hiDAs) were infected with scAAV2/retro vectors encoding scFv-833D4, scFv-73B3, or a control scFv, and subsequently exposed to 488-hPFFs. Two hours post-exposure, cells were washed, fixed, and the Alexa Fluor 488 intensity within TH^+^ dopaminergic neurons was measured. We found that expression of either scFv-833D4 or scFv-73B3 significantly reduced the 488-hPFF fluorescence intensity inside dopaminergic neurons compared to cells exposed to the control scFv (**Figure 7E-F**). This suggests that both scFvs effectively inhibit the entry of exogenous aSyn PFFs into human neurons.

### Systemic delivery of scAAVs encoding anti-aSyn scFvs do not induce an inflammatory response in mice

Since inflammation and auto-immune responses are important concerns with immunotherapy for neurodegenerative disorders, we wished to investigate if the systemic injection of scAAV-scFv induced an inflammatory reaction. We used transgenic mice that express a dual bicistronic reporter system of GFP and Luciferase under the transcriptional control of the murine Toll-like receptor 2 (TLR2) gene promoter [57]. Transcriptional activation of TLR2 is indicative of microglial activation and has been associated with aSyn pathology [58,59]. We performed bioluminescence imaging at days 0, 4, 14 and 28 after IV injection of AAV-scFv-control, AAV-scFv-833D4, and AAV-scFv-73B3. Additionally, an AAV encoding GFP served as a non-self protein comparator, saline was used as a negative control, and intraperitoneal injection of lipopolysaccharide (LPS) was used as a positive control. Four days post-injection, a strong bioluminescence signal was observed in the LPS group, while the other groups displayed only slight increases in signal compared to the saline group (**Figure 8A-C**). Furthermore, we wished to evaluate if the scAAVs encoding our anti-aSyn scFvs caused a long-term increase in pro-inflammatory cytokines indicative of an immune response. To do so, we used mice that received intrastriatal PFF (or PBS) with or without intravenous scAAV-scFv administration (833D4, 73B3, control or saline). We employed a mouse cytokine array that allowed us to visualize the level of 70 cytokines in the brain of the mice at the time of their sacrifice (maximum 12 weeks post-PFFs). Analysis of the relative dot intensity for each cytokine did not reveal significant differences in the level of cytokines most frequently found elevated in PD patients [60] such as TNF-α (**Figure 8D**), INF-γ (**Figure 8D**), IL-1β (**Figure 8F**) or IL-10 (**Figure 8G**) between the mice that received the scAAV-scFvs compared to uninjected mice. We did observe an increased IL-12 concentration in PFF-injected mice compared to PBS-injected (**Figure 8H**), although there were no differences between the different AAV-scFv groups. IL-12 is a pro-inflammatory cytokine that is secreted by various immune cells, including microglia [61]. These results indicate that the systemic injection of scAAV encoding anti-aSyn scFvs does not cause sizable inflammation in the brain of the mice and does not drastically modify the long-term immune response following PFFs injection.

**Figure 8:**
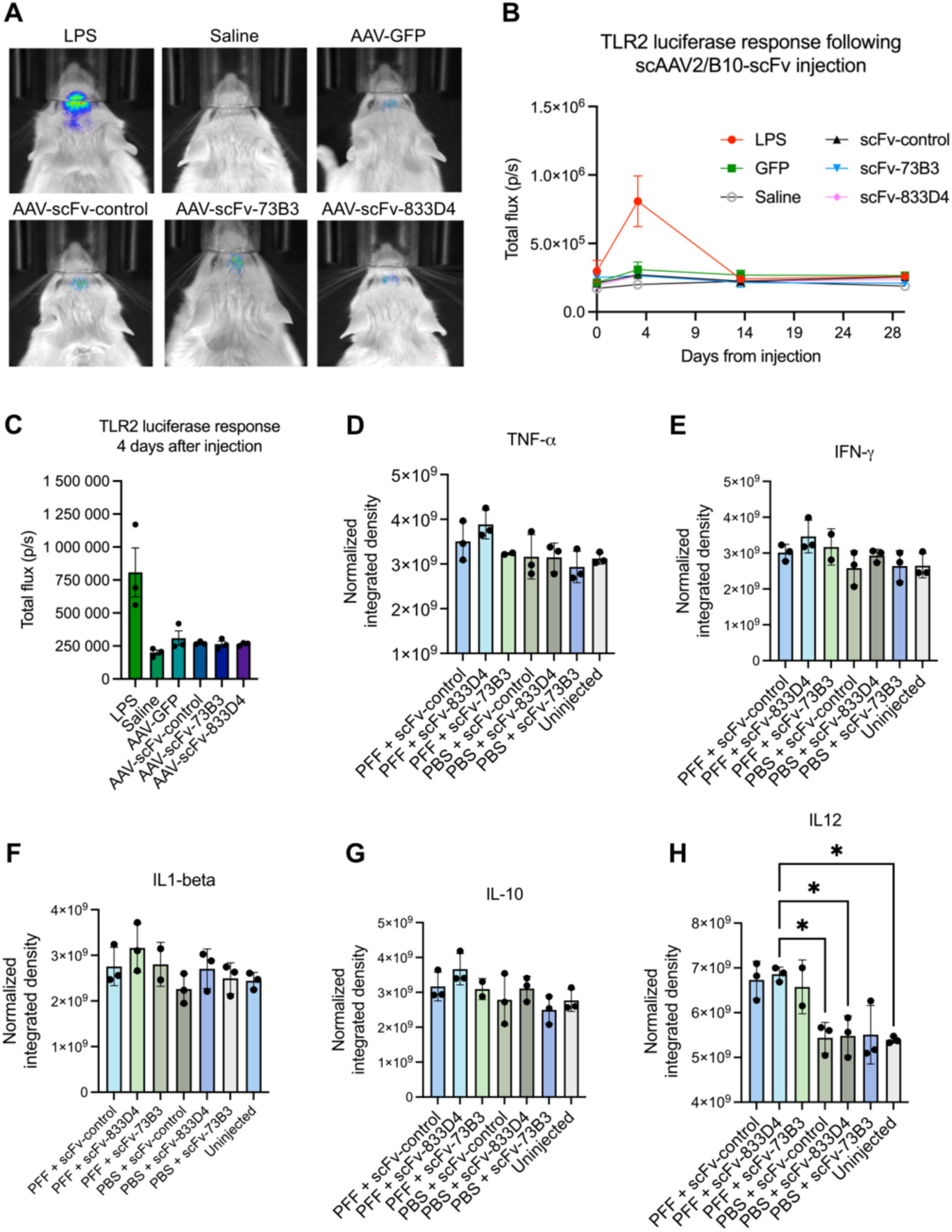
Systemic injection of scAAVs encoding anti-aSyn scFvs does not induce an inflammatory response in mice. A-C) Mice expressing luciferase under the control of the TLR2 promoter were used to evaluate the temporal brain inflammation caused by IV injection of the scAAV-scFvs. A) Representative images of the bioluminescence signal at 4 days post injection. B) Total flux (in photon/second), corresponding to the level of induction of TLR2 expression as a function of the time from injection. C) Total flux at day 4 post-injection showing the absence of TLR2 induction following the injection of the scAAV-scFvs. D) Normalized integrated density of TNF-α (D), IFN-γ (E), IL-1β (F), IL-10 (G) and IL-12 (H) in brain lysates 12 weeks post-PFFs injection (n=3/group) measured with a cytokine array. One-way ANOVA with Dunnett’s multiple comparison test *p<0.05. G-I). G). H) I)

## Discussion

Parkinson’s disease and other Lewy body diseases are affecting a growing number of people worldwide. Treatment options are currently solely symptomatic, with no modification of disease progression [62]. Patients are in desperate need of better therapeutics to ameliorate their quality of life. Immunotherapy is extensively studied for neurodegenerative disorders, including Alzheimer’s (AD) and Huntington’s disease, but symptomatic improvement has remained limited so far in clinical trials [63]. Although target engagement is generally good, the choice of the target itself and the off-target/secondary effects have been major obstacles in the field [64]. Because antibodies can recognize specifically a mutated, post-translationally modified or even aberrantly folded protein, they are potent tools to address proteinopathies. Monoclonal antibodies targeting aSyn have shown promising results in PD models and clinical trials [24–26,30,31,65,66]. However, immunoglobulins (e.g. IgGs) administered intravenously can only cross the blood-brain barrier inefficiently, typically 0.1% of the injected mass [67] and have a half-life between 15 and 30 days [68]. Repetitive administration of high doses of an antibody would thus be necessary to achieve long-term beneficial effects in the brain, which reduces patient compliance and increases treatment costs. To get around both the difficult access to CNS and the phasic nature of traditional antibody administration, achieving long-term, tonic production of antibodies or derivatives through CNS somatic gene transfer is an attractive avenue.

We generated novel mouse monoclonal antibodies to p129-aSyn and pan-aSyn. Among the antibodies obtained, we selected 4 IgGs that showed specific and strong signal towards pan- or phosphorylated aSyn in immunofluorescence and western blot (**Figure 1**). Our newly generated antibodies could bind to aSyn inclusions in brain sections from patients suffering from PD and DLB (**Figure 2**), which suggests that they could bind *in vivo* to endogenous aSyn and have a potential therapeutic capacity. The weaker binding to MSA aggregates suggests a potential conformational specificity, as it was shown that MSA pathology is caused by a different aSyn strain [69]. The IgG 833D4 could prevent fibrillization of aSyn in a Thioflavin T aggregation assay, which indicates a possible inhibitory action on aSyn aggregation. We generated secreted single-chain variable fragments (scFvs) from IgGs 833D4 and 73B3. In our synucleinopathy model, intravenous injection of scAAVs encoding tagged scFvs resulted in a widespread distribution of scFv-producing brain cells with little if any scFv expression in the liver, the main off-target organ of human AAVs. The brain expression of scFv-833D4 and 73B3 prevented motor impairments and significantly decreased the number of pS129-aSyn inclusions in the brain of the mice (**Figure 5-6**). We confirmed that the scFvs can be internalized by neurons and microglia using a bicistronic construct allowing the expression of both a secreted scFv and a non-secreted GFP from the same promoter. This suggests that the scFvs can act on intracellular aSyn (**Figure 7**). Moreover, infection of hiDAs with scAAVs encoding our secreted scFvs could prevent the internalization of 488-hPFFs (**Figure 7**) pointing to a modulation of aSyn pathology transmission. It is thus possible that the anti-aSyn scFvs can have a dual effect on intra- and extracellular aSyn. In mice, the intravenous injection of scAAVs encoding our anti-aSyn scFvs did not provoke brain inflammation (**Figure 8**), a concern for immunotherapy targeting proteins from the self [70]. The absence of an Fc region in scFvs probably explains this advantage, as IgGs interact and activate immune effector cells and molecules through their Fc region. Thus, the mechanism of action of our scFvs is thought to be a neutralizing action that inhibits transmission and aggregation of aSyn. Further experiments would be needed to elucidate the exact neuroprotection mechanism.

In Lewy body diseases, the chronological process of aSyn dysregulation and subsequent cell failure is not well understood. Phosphorylation of aSyn at the serine 129 has been considered a pathological hallmark for several years because of its abundance in the brain of PD patients and its very low amount in healthy brains [71]. Levels of pS129-aSyn have also been positively correlated with disease severity in humans [72]. However, conflicting results have been obtained regarding the effect of pS129 on aSyn aggregation and transmission [73,74]. Because of this uncertainty, we decided to evaluate the therapeutic potential of a pan-aSyn specific scFv (833D4), and one that recognizes preferentially pS129-aSyn (73B3). Targeting the phosphorylated protein would avoid potential secondary effects caused by down-regulation and loss of function of the normal protein [75,76]. However, with the dose of scAAV-scFv used, and possibly because of the fact that the scFvs are secreted, we did not observe a reduction in total aSyn protein level in our model (**Figure 6**). We believe it is possible to target total aSyn and to keep physiological subthreshold protein level. Moreover, several studies have suggested that phosphorylation is a late event in aSyn dysregulation and that it likely follows aggregation [50,77,78]. It could thus be advantageous to target the non-phosphorylated form before extensive aggregation.

With our SPR experiment, we surprisingly found that the antibody 73B3 has a relatively high binding affinity to non-pS129-aSyn, which was not observed in any other experimental setting. KD values alone do not provide information about the binding kinetics to the antigen. By looking at the response curve, we can observe that the dissociation rate of 73B3 is much faster with aSyn compared to pS129-aSyn (**Figure 3**). This correlates with an affinity of the 73B3 that is 5 times higher towards pS129-aSyn and explains the preferential binding to this species over the non-phospho protein. When using the IgG 73B3 in an immunoassay, it is likely that the washing steps almost completely eliminate the binding to the non-phospho aSyn. To our knowledge, none of the commercially available antibodies claiming to be specific for pS129-aSyn have published SPR data or equilibrium dissociation constant (KD) values for both the phosphorylated and non-phosphorylated forms of the protein. Interestingly, the 833D4 antibody showed higher affinity towards both aSyn and pS129-aSyn compared to the 73B3 antibody (**Figure 3**). This could explain the slightly better efficiency of the scFv-833D4 in the behaviour experiments and the histopathological analyzes (**Figure 5 and 6**).

Our approach is unique in several aspects compared to other studies using antibodies or antibody derivatives for PD treatment [79]. First, our therapeutic is administered non-invasively by an intravenous injection. Brain bioavailability has always been a challenge for brain-targeted therapeutics. Injecting AAV-encoded scFvs directly in the brain is possible [80] but involves high surgery-related risks and reduces treatment accessibility. We believe that systemic delivery is ideal and we achieved it in our study with the AAVCap-B10 vector, which easily penetrates the blood-brain barrier in mice and marmosets [55]. We used a generic CMV promoter/enhancer to achieve high and rapid brain expression and observed minimal expression in the liver, consistent with the previously described properties of the B10 capsid [55]. The use of a brain-specific and/or inducible promoter could be favored for human trials as it reduces the risks of adverse immune reactions against the transgene. If proven safe, whole-body AAV-scFv expression could however provide the advantage of acting on peripheral Lewy body pathology [19]. Studies on non-human primates would be needed to evaluate safety and pharmacokinetics. Secondly, in contrast with several neuroprotection studies for PD [25,26,80,81], we administered our virally encoded anti-aSyn scFvs after the induction of the pathology in our model. Although we did not monitor the time course of the scAAV transduction in our model, we hypothesize based on previous reports that the expression of the scAAV and the concentration of scFv in the brain is maximal 1 to 2 weeks after injection [82–84]. This likely means that PFFs internalization and subsequent seeding has already begun when the treatment is started [85,86]. This makes our approach more translational, as dopaminergic degeneration is already prominent when most of the patients are diagnosed [87]. Even by administrating the scAAV-scFvs after the disease onset, we could obtain a good protection against aSyn neuropathology. It would nevertheless be interesting to inject the scAAV-scFvs at different time points to evaluate if the pathology can be stopped or even reversed at a later time. Thirdly, the targeting of extracellular aSyn distinguishes our strategy from others as most of the previously described antibody derivatives against aSyn were intrabodies, that are solely expressed intracellularly [79]. By keeping the signal peptide from the parental IgGs of our scFvs, we allow their secretion in the extracellular space. This has two main advantages: 1) We can act on cell-to-cell transmission by targeting extracellular aSyn, and 2) We can potentially obtain a better therapeutic effect as cells that are not directly infected by the scAAV can be reached by the scFvs. Results from a previous study suggested that secreted scFvs can penetrate cells and act intracellularly [52], and we replicated these observations with our AAV-scFv-IRES-GFP construct. It is thus possible that our anti-aSyn scFvs also act on intracellular aSyn, possibly inhibiting aggregation. Hence, monomeric or small oligomers of aSyn could potentially be cleared more easily by intracellular degradation mechanisms (e.g., proteasome).

M83 transgenic mice with intrastriatal aSyn PFFs injection represent an attractive and popular synucleinopathy model due to the rapid but progressive development of Lewy body-like aggregates in various brain regions. This neuropathology is also accompanied by severe motor symptoms. However, the absence of dopaminergic degeneration and lack of aSyn pathology in TH+ neurons of the SNc make it an imperfect sporadic PD model [53,88]. Furthermore, high intra-group variability in the motor phenotype and histopathology has been observed in our study and is possibly due to two main factors: the variable level of transgene expression in M83^+/-^ mice, and the intrinsic variability of recombinant aSyn PFFs. M83 mice have been generated by random insertion of the transgene and are maintained on a mixed genetic background B6:C3 [53]. The random chromosomal insertion and the mixed background can influence the expression level of the transgene and cause inter-individual heterogeneity [89–92]. This could affect the seeding efficiency and the cell-to-cell transmission of the injected PFFs and result in a nonuniform phenotype. Additionally, the properties of recombinant aSyn PFFs have been shown to fluctuate from batch to batch and to induce varying degrees of pathology when injected intracerebrally [93]. Overall, the combined properties of the transgenic mice and the recombinant PFFs likely explains most the variability in the motor phenotype and the neuropathology obtained. To avoid part of this unpredictability, researchers should ensure that all animals are injected with the same batch of PFFs when using this model.

In summary, we designed virally encoded scFvs against aSyn as a therapeutic strategy for Lewy body diseases. Injection of scAAVs encoding scFv-833D4 and scFv-73B3 prevented motor impairments and aSyn neuropathology in a synucleinopathy mouse model, without causing neuroinflammation. Our approach is minimally invasive and shows a long-term efficiency. We believe our strategy could be adapted for PD and other Lewy body diseases patients to provide neuroprotection and delay disease progression.

## Methods

### Cell culture

HEK cells were maintained in DMEM (Life Technologies #11995065) supplemented with 10% fetal bovine serum (Life Technologies #16000036) and 1% penicillin-streptomycin (Cytiva #SV30010). For immunocytochemistry, cells were plated on Poly-L-lysine (Sigma #P4707) - coated glass coverslips (Fisher #12-545-81) at a density of 50K/well. The following day, cells were transfected with 500ng DNA/well using Lipofectamine2000 and incubated for 48h at 37°C. Cells were fixed in 4% PFA on ice for 20min and washed with PBS afterwards. For western blots experiments, cells were plated in 6-well plates at a density of 400K cells/well and transfected the next day with 3ug total DNA/well using Lipofectamine2000. 48h later, cells were lysed with RIPA buffer (NEB #9806S) supplemented with PMSF on ice for 5 minutes. Lysates were transferred to 1,5mL tubes, sonicated briefly, and centrifuged at 14000g for 20minutes at 4°C. Supernatant was transferred to a clean tube and total protein concentration was measured with the DC protein assay (Bio-Rad #500-0112). 4X Laemmli Sample buffer (Bio-Rad #1610747) was added at the appropriate concentration and the lysates were boiled 5min at 95°C. Final lysates were aliquoted in and stored at -80°C.

### Human induced pluripotent stem cells (hiPSCs)-derived dopaminergic neurons

For the 488-PFF experiment in hiPSCs-derived dopaminergic neurons, the line AST23 was used. The cells were sourced from the C-BIG repository (The Neuro, Montreal, Canada). These cells have a triplication of the *SNCA* gene, resulting in 4 copies of *SNCA* and were described in [56]. hiPSCs were maintained in mTESR-Plus medium (Stemcell, 100-0272) in 6 well plates coated with Matrigel (Fisher, CB-40230C) and were passaged every other day using Accutase (Stemcell, 07920) or Gentle Cell Dissociating Reagent (GCDR) (Stemcell, 7174). Accutase was used to obtain a single cell suspension before starting a differentiation. Briefly, 0.5ml of Accutase was added per well and the plate was incubated 3-6 minutes at 37°C. When cells started to detach, they were resuspended in KO-DMEM (Life Technologies #10829-018), centrifuged 5min at 300xg, and resuspended in mTESR-Plus medium. GCDR was used to obtain cell aggregates. Briefly, 1ml of GCDR was added per well, and the plate was incubated 8-10 minutes at room temperature. The GCDR was then removed and replaced with room temperature mTESR. The cell colonies were gently detached with a cell scraper and transferred to a 15ml tube. The cell mixture was pipetted up and down to create a uniform suspension of aggregates of about 50-200μm, and then plated in mTESR-Plus at the desired density. Rock inhibitor (Stemcell, 72304) (10 uM) was added to the media upon passaging. Differentiation to dopaminergic neurons was carried as previously described in [94]. At DIV 35, cells were plated on 12mm glass coverslips at a density of 1.2 million cells/ml. Infection with AAV2/retro encoding each scFv was carried at DIV 38 and 43 with 7.5E9 GC/well. Five days after the second infection, 1.2 ug/well of 488-PFF was added to the cell media for 2 hours. Cells were washed 2 times with cold PBS before fixation with 4% PFA on ice.

### Monoclonal antibody generation and testing

This service was provided by MédiMabs (141 avenue du Président-Kennedy, Montréal, Québec, Canada). Six mice were immunized 3 times with a 25aa peptide corresponding to a region near the C-terminal part of aSyn with the serine 129 phosphorylated (See Figure 3A). The peptide was coupled to KLH (Keyhole Limpet Hemocyanin) for its immunogenic effect. Splenocytes were harvested from 3 mice that developed the strongest positive immune response (antibody titers were measured by ELISA) and were fused to NS0 myeloma cells to generate hybridomas. Hybridomas were plated at 10, 1 and 0.1 cell well, and supernatants from clonal hybridomas was tested by ELISA against the peptide crosslinked to BSA.

Hybridoma supernatant from ELISA-positive clones was used as a primary antibody solution in western blot and immunocytochemistry (ICC) to verify specificity against phosphorylated versus total aSyn protein. For ICC, HEK cells were plated on coverslips and transfected with either aSyn, aSyn + PLK2, aSyn-S129G + PLK2 or GFP as described before. After fixation, cells were incubated with a blocking buffer (1X PBS + 5% normal donkey serum + 1,5% TritonX-100) for 1h at RT. Hybridoma supernatants were used as primary antibodies O/N at 4°C. The next day, cells were washed with PBS, and secondary antibody (AlexaFluor647 anti-mouse) was added in blocking buffer for 1h at RT. Cells were mounted with DAPI-containing mounting media (Cat# P36935). For WB, 20ug DNA was loaded in a pre-casted SDS-PAGE gel (Bio-Rad #4568093) and the gel was run for 90min at 100V. Proteins were transferred on a nitrocellulose membrane using the TransBlot Turbo apparatus (Bio-Rad). Membranes were blocked for 1h in TBST containing 5% skim milk and then incubated O/N at 4°C in the hybridoma supernatants. The next day, membranes were washed in TBST and incubated 1h at RT in 5% milk with the secondary antibody (HRP-anti-mouse). Images were acquired with Bio-Rad’s Chemidoc using ECL substrates (Cat# 1705062).

### Monoclonal antibody purification

Monoclonal antibodies were purified from the supernatant of hybridoma cultivated in a medium containing extra-low IgG serum (Gibco). Briefly, Protein-G-Agarose resin was added to a 50mL tube and washed 2 times with TBS. Hybridoma supernatant was added to the tube and incubated overnight with slow agitation at 4°C. The next day, the tube was centrifuged, and the supernatant was discarded. The resin was then washed 3 times with TBS and a glycine solution (pH 3.5) was added to detach the antibody from the resin. After a 5 minutes incubation, the tube was centrifuged, and the supernatant was collected and placed in another tube containing 1M Tris. The purified antibody was then concentrated with a Vivaspin 6 column with a 10kDa molecular weight cut off membrane. Concentration was read with a nanodrop system (Thermo Scientific).

### Construction of scFv-encoding scAAV2 plasmid vectors

The entire sequences of the light and heavy chains of the selected IgG antibodies were sequenced using NGS by MédiMabs (Montréal, Canada). scFv-encoding ORFs were designed in silico, synthetized into double-strand DNA using oligonucleotide PCR assembly (by Bio-Basic, Canada), and cloned into plasmid Bluescript-SK-II (StratageneTM) as BamHI-NotI fragments. The anti-aSyn scFv open reading frames generated encode (5’ to 3’) the IgG light chain variable domain with its signal sequence, followed by a synthetic sequence encoding a (Gly_4_Ser)_4_ flexible linker and the Heavy chain variable domain, followed in frame with a HA tag and a Myc tag and terminal stop codon. In some cases, codon wobbling was used to get rid of other BamHI or NotI restriction sites within the ORFs. All sequences were verified by Sanger sequencing. The synthetic scFv-encoding genes were then subcloned in an scAAV cloning vector plasmid derived from pscAAV-CMV-EGFP (a kind gift from J. Samulski, Gene Therapy Center, University of North Carolina at Chapel Hill, USA [84]). The cloning vector plasmid was modified so as to contain within the two ITRs the CMV promoter/enhancer sequence from CMV-CHFR-mGFP (GenBank: MK317917.1) followed by the short SV40 intron, the BamHI and NotI restriction sites for cloning, and an optimized combined WPRE-sv40 polyadenylation signal we termed WpPA [51]. Plasmids were transformed in 5’-Alpha or Stable competent bacteria (New England Biolabs, Cat C3040H and Cat C2987H), and extracted using a commercial kit (Invitrogen^TM^ K182002). Constructs sequences were verified by digestion with restriction enzymes, and by Sanger sequencing. The scAAV-control contains an expression cassette for an anti-GFP scFv (Restriction enzymes were from New England Biolabs Canada. AAV2/retro and AAV2/B10 viral vectors were produced with the scFv-encoding plasmids (pscFvs) by the Molecular Tools Platform at CERVO Research Center.

### Generation of scAAV2/CapB10 viral particles encoding scFvs

scAAVs were produced by the Canadian Neurophotonic Platform Viral Vector Core (Quebec city, Canada). Briefly, self-complementary AAV2 viral vectors encoding secreted scFvs were generated by co-transfection of either the scAAV-scFv-833D4, scAAV-scFv-73B3 or scAAV-scFv-control (encoding a secreted anti-GFP [52]) with helper plasmid pXX680, a plasmid containing the AAV2/CapB10 Rep/Cap in 293T17 cells. 48h post transfection, the cells were harvested by gentle scraping and were pelleted by centrifugation. Viral particles were released by 4 freeze-thaw cycles on dry-ice/ethanol and free DNA was digested with Benzonase. AAV particles were purified by a discontinuous gradient of iodixanol (15-25-40-60%) and ultracentrifugation (70,000 RPM 1h30 at 16°C). The viral preparation was then concentrated through a Centricon filter and adjusted to 5% sorbitol and 0.001% Pluronic acid for storage and experimentation. Titration was performed by digital droplet PCR using primers specific for AAV2 ITRs.

### Dot blots

aSyn monomers (Proteos #RP-003), pS129-aSyn monomers (Proteos #RP-004) and BSA (Sigma #A9647) were manually blotted on nitrocellulose membranes. Each dots contains 2ug of total protein in 2ul volume. aSyn and pS129-aSyn were diluted with BSA solution to achieve constant protein amount. Membranes were let to dry for 15min a RT and were then humidified with 1X TBS. Membranes were blocked with TBST containing 5% skim milk for 1h at RT. HEK cells plated in 6-well plates were transfected with the pscFvs and the cell media was collected 2 days later. The medium containing the secreted scFv was used as a primary antibody for O/N at 4°C. The next day, membranes were incubated with an HRP-conjugated myc antibody for 1h at room temperature. Images were acquired with Bio-Rad’s Chemidoc using ECL substrates (Cat #1705062).

### ELISA

ELISA high binding plates were coated by incubating with a solution of purified alpha-synuclein (20ug/mL) in carbonate binding buffer overnight at 4°C. The next day, the plate was washed 3 times with wash buffer (1X PBS containing 0,05% Tween 20) and then incubated for 1h at room temperature in blocking buffer (3% fish gelatin in PBS). After blocking, the plate was washed, and primary antibody diluted in 1% fish gelatin in PBS or filtered scFv-containing cell media was added and incubated for 2h at room temperature. The plate was washed again and incubated for 30min at 37°C with secondary antibody conjugated to HRP (see supplementary material). After washing, 100μl of Tetramethylbenzidine solution (Sigma T4444) was added and incubated at 37°C for 5-10 minutes. 100μl of 1M HCl was added to each well to stop the reaction and the absorbance at 450nm was read with the Victor Nivo plate reader (PerkinElmer). Experiment was repeated two individual times.

### Surface plasmon resonance

Surface Plasmon Resonnance was performed by Proteogenix (19 rue de la Haye, Schiltigheim, France). The Biocore^TM^T200 was used for SPR analysis. Briefly, aSyn or pS129-aSyn monomers (Proteos #RP-003 or #RP-004) diluted in 10 mM NaAc (pH 5.5) were immobilized on a CM5 sensor chip using maleimide EDC/NHS coupling. Antibodies were flown over the CM5 chip at concentrations ranging from 0.5 to 16 nM and the response was captured over time. The running buffer and the antibody dilution buffer was HBS-T (50 mM Hepes-NaOH pH 7.4, 150 mM NaCl and 0.005% Tween-20). Regeneration with glycine pH 1.5 was performed between each concentration tested. The kinetics parameters and affinity were calculated using BIAevaluation software.

### Thioflavin T aggregation assay

aSyn monomers (Proteos #RP-003) or pS129-aSyn monomers (Proteos #RP-004) at 2mg/ml were incubated with 250μg/ml of IgG 833D4, 73B3 or unspecific control (IgG2a against LaCrosse virus glycoprotein G, ATCC CRL-2290). Volume was completed to 100μl with 2X PBS. Tubes were sealed with parafilm and incubated for 7 days in a Thermomixer at 37°C with 1000 RPM agitation. At day 3, 5 and 7, an aliquot of 5μl was taken (in duplicate) and mixed with 25μM Thioflavin T in a black non-binding 96 well plate with flat bottom. The plate was incubated 15 minutes at room temperature and absorbance was read with the Victor Nivo^TM^ plate reader (PerkinElmer) (ex = 450nm, em = 480 nm). Experiment was repeated two individual times.

### Mice

A53T a-synuclein transgenic line M83 (B6;C3-Tg(Prnp-SNCA*A53T)83Vle/J, JAX stock #004479) was used. Mice were breed in house by crossing homozygous females with hemizygous males. Pups were genotyped using ddPCR according to Jackson laboratory’s protocol. Hemizygous females aged between 60 and 90 days were used. Mice were housed in a light-dark standard cycle with access to water and food ad libitum and Enviro-dri (ScottPharma Solutions Inc.) was added to their cage to encourage nesting. All mice were weighted weekly starting at 8 weeks post-PFF injection. Animal protocol was approved by the Laval University Animal Protection Committee (CPAUL) and by the Canadian Council of Animal Care (CCAC).

### PFF preparation

Human aSyn PFF were provided by Thomas Durcan’s laboratory, at McGill University (Montreal, Canada). They were prepared according to standard operating procedures based on the protocol described in [95]. The protocol for the production of the PFF can be found here: https://zenodo.org/record/3738335#.Yp46fXbMKUl. The protocol for characterization can be found here: https://zenodo.org/record/3738340#.YzIjBezML0p.

### In vivo injections

Mice were injected with aSyn pre-formed fibrils (PFF) (kindly provided by Thomas Durcan’s lab, McGill University) in the right dorsal striatum (coordinates: A/P +0.20; M/L + 2.00; D/V 2.6) under isoflurane anesthesia. The needle was left in place for 5 minutes after injection, and then slowly pulled back. Before injections, PFFs were sonicated in a Bioruptor Pico bath for 20min (30sec. ON, 30sec. OFF, 20 cycles) at 10°C. PFFs were injected less than 4h after sonication to avoid degradation. A total of 5μg (2.5mg/ml) of PFFs in a volume of 2μl was injected.

7 days after PFF injection, mice were injected intravenously in the retro-orbital sinus with either one of the scAAV2/CapB10-encoding an scFv (either scAA2/capB10-scFv833D4, scAA2/capB10-scFv73B3, or scAAV2/CapB10-scFv-control). Briefly, mice were anesthetized with isoflurane and a solution of alcaine was applied on the eye for 30 seconds. Insulin syringes were used to inject 100μl of scAAV diluted in Ringer’s lactate. 5E11 GCs of virus was injected per mouse.

### Behavioral tests

#### Clinical scoring

Starting 8 weeks post-injection, mice were monitored for motor impairments using the grid presented in supplementary. 3 aspects were evaluated: clasping, gait, and ledge walking. Each test can be scored from 0 to 3 for a maximum of 9 points for all tests. See supplementary material.

#### Rotarod

Mice were placed in the behavior room 1h prior to the test for habituation. For 2 days before the test. Mice were trained by placing them on the rod (IITC Life Science) for 5min at 4RPM constant speed. The third day, mice were tested in 3 trials of 5min with speed increasing from 4 to 40 RPM. 5 minutes breaks were allowed between trials where the mice remained in the apparatus.

#### Grip strength test

Mice were placed in the behavior room for 1h prior to the test for habituation. The grip strength test from Biosed company was used (#EB1-BIO-GS4). Mice were held by the tail and were lowered on the grid (#EB1-BIO-GRIPGS) until their 4 paws gripped and were then pulled with constant strength horizontally until the hindlimbs let go. Maximum strength displayed was noted and 6 trials were made. Maximal and minimal values were excluded from analysis and a mean of the 4 remaining values was normalized on the mice weight.

### Tissue processing

For histopathological analysis, mice were anesthetized with ketamine-xylazine and were sacrificed by cardiac perfusion with cold PBS followed with 4% paraformaldehyde. Brains and spinal cords were extracted and post-fixed for 90min in 4% PFA, then transferred in 30% sucrose solution for O/N at 4°C. Brains and spinal cords were frozen in OCT and cut coronally at 60um with a cryostat (Leica). Ipsilateral cortex was marked with a blade to recognized injected side. For western blot analysis, mice were sacrificed by cervical dislocation without anesthesia. Brains were extracted and cortex, hippocampus, midbrain, striatum, and medulla were dissected and frozen quickly in liquid nitrogen. The tissue was stored at -80°C until used. Upon thawing on ice, tissue was lysed manually with a potter in RIPA buffer (NEB #9806S) supplemented with PMSF (NEB #8553S). After quick sonication, lysates were centrifuged at 14000g for 20 min at 4C. Supernatant was transferred to a clean tube and total protein concentration was measured with the DC protein assay II (Bio-Rad #500-0112). Samples were stored at -80 until used.

### Western blotting

4X Laemmli sample buffer (Bio-Rad #1610747) was added, and samples were boiled for 5min at 95°C and then loaded on 4–20% Mini-PROTEAN® TGX Stain-Free™ Protein Gels (Bio-Rad #4568093). Gel was run at 100V for 90 minutes and proteins were transferred to a nitrocellulose membrane using the Bio-Rad Transblot Turbo system. For pS129-aSyn staining, membranes were fixed for 15min in 4% PFA and then washed 3 times 10min with 1X TBS. 5% milk in TBST was used as a blocking solution for 1h at room temperature. Membranes were incubated with primary antibodies overnight at 4°C in 5% milk (Supplementary table 3). After washing in TBST, membranes were incubated in secondary antibody solution in 5% milk for 1h at RT. Images were acquired with Bio-Rad’s Chemidoc using ECL substrates (Cat #1705062). Analysis was performed with ImageLab.

### Immunohistochemistry

Antigen retrieval was used for NeuN staining only. Briefly, brain sections were incubated in pre-heated sodium citrate buffer at 95-100°C for 15min and then washed 2 times in PBS with 0,05% TritonX-100. Sections were washed in 1X PBS and then incubated in blocking solution (1X PBS + 5% NDS + 0,3% TritonX-100 or 1X PBS + 5% BSA + 0,3% TritonX-100) for 1h at RT. Sections were incubated O/N with primary antibodies in blocking solution (See Supplementary table 3). The next day, sections were incubated with secondary antibodies (AlexaFluor 555 [A21436]; AlexaFluor 647 [A31571]; AlexaFluor 488 [A21206]; Dylight 405 [713-475-147]) in blocking solution for 1h at room temperature. If Amytracker (Amytracker 520 Ebba Biotech) was used, it was incubated for 30 minutes at a dilution of 1:1000 in PBS before mounting. Images were acquired with the Zeiss LSM-710 confocal microscope and analyzed with ImageJ.

### Myc staining

Staining of human Myc tag in brain and spinal cord sections was done using either the 9E10 purified Mab (Invitrogen #132500), or 9E10 hybridoma supernatant as primary antibodies, with the Alexa Fluor 555 Tyramide Superboost kit (Invitrogen #B40923) according to the manufacturer’s instructions. Anti-HA immunostaining was performed using Abcam ab9110 primary antibody.

### Cytokine array

The Mouse Cytokine Array C3 from RayBiotech (AAM-CYT3-8) was used according to manufacturer’s instructions. Brain lysates from the medulla and midbrain were mixed equally for a total protein amount of 250μg. Images were acquired with Bio-Rad’s Chemidoc. Analysis was performed with ImageJ according to the manufacturer’s instructions.

### Cell counting

For TH and NeuN cell counting, a stereological counting software was used (StereoInvestigator Offline, MBF Bioscience). 4 midbrain sections (spaced by 180 μm) were analyzed per animal. A surface of 55% of the SNc was counted manually in random places for the software to estimate the total cell number.

### Image analysis

pS129-aSyn was quantified in 4 brain regions (midbrain, PAG, pons and medulla). Two consecutive brain sections (of 60 μm) were analyzed per mouse. An automatic threshold was set for all images and the particle analysis of Fiji was used to quantify the area. For analysis of TH and DAT in the striatum, corrected total cell fluorescence was used (CTCF = Integrated density – (Area of selected region x Mean fluorescence of background readings)). A ROI corresponding to the dorsal striatum was traced and measured on 4 sections for each animal. Mean values are reported as ipsilateral/contralateral fractions.

### Statistical analysis

All statistical analyzes were performed using GraphPad 9.4.0. Statistical significance was determined by unpaired two-tailed Student’s *t* test for 2 groups comparisons, or one-way ANOVA test with Tukey’s or Dunnett’s correction for multiple groups comparisons. *P* values lower than 0.05 were considered significant. \**P*<0.05, \*\**P*<0.01, \*\*\**P*<0.001, \*\*\*\**P*<0.0001.

Experimenters were not blind to the groups when performing analyses.

## Data availability

The data that support the findings of this study are available from the corresponding author, M.L., upon reasonable request.

## Acknowledgements

We thank Véronique Rioux for the technical and administrative assistance and the Brain Bank of the CERVO Research Center for the human brains samples. We thank the Canadian Neurophotonic Platform Viral Vector Core for the scAAV viral vector production. This work was supported by a Michael J. Fox Foundation grant (pre-clinical program). A-M.C. was supported by a doctoral fellowship of the Canadian Institutes of Health Research (CIHR). B.M received a master’s scholarship from Fonds de recherche du Quebec – Sante (FRQS). M.L. received salary support from FRQS in partnership with Parkinson Quebec.

## Author contributions

A-M.C contributed to conceptualization, methodology, experimentation, analysis, writing and editing. B.M. contributed to some experiments and related analysis. M.P. provided the human brain sections. T.D. laboratory provided the human aSyn PFF for in vivo and in vitro experiments. W.L and I.S. contributed to the production and characterization of human aSyn PFF. J.P.J. contributed to conceptualization. C.G. contributed to conceptualization, methodology, experimentation, manuscript revision, supervision, and funding acquisition. M.L. contributed to conceptualization, methodology, writing, supervision, project administration and funding acquisition.

