## Supplementary material for "Virally encoded single-chain antibody fragments targeting alpha-synuclein protect against motor impairments and neuropathology in a mouse model of synucleinopathy"

#### *Human brain staining information:*

**Supplementary table 1**: Information about the patients for each pathological brain analyzed.

| Brain ID | Age | Sex | Post-mortem delay (h) | Diagnostic | Disease duration (years) | Pathologist notes |
|---|---|---|---|---|---|---|
| C-002 | 78 | M | 2.5 | PD | 9 | Some Lewy bodies in frontal lobe |
| C-004 | 73 | F | 18 | PD | 6 | Some Lewy bodies in frontal lobe |
| C-097 | 55 | M | 31 | PD | 14 | Lewy bodies in frontal lobe |
| C-0111 | 64 | F | 19.5 | MSA | 13 | MSA-C: Some inclusions in oligodendrocytes in the cortex |
| C-0081 | 49 | F | 19 | MSA | 4 | MSA-P: Inclusions in oligodendrocytes in the cortex |
| C-0030 | 66 | M | 11 | MSA | 12 | MSA-P: Inclusions in oligodendrocytes in the cortex |
| C-0024 | 78 | M | 15 | DLB | N/A | Lewy bodies in frontal lobe |
| C-0031 | 74 | M | 5 | DLB | 19 | Lewy bodies abundant in frontal lobe |
| C-0080 | 84 | M | 16.5 | DLB | 5 | Lewy bodies in frontal lobe |

**Supplementary table 2**: Relative abundance of aggregate staining in each brain analyzed.

| | | | Level of aggregates staining | | | | | |
|---|---|---|---|---|---|---|---|---|
| **Brain ID** | **Region** | **Condition** | **73B3** | **73D1** | **833D4** | **826B2** | **CST2628** | **BioLegend** |
| C-0002 | SN | PD | +++ | ++ | ++ | ++ | N/A | N/A |
| C-0004 | SN | PD | +++ | ++ | ++ | ++ | N/A | N/A |
| C-0097 | SN | PD | +++ | +++ | + | ++ | + | +++ |
| H-0172 | SN | CTRL | - | - | - | - | N/A | N/A |
| C-0129 | SN | CTRL | - | - | - | - | - | - |
| C-0111 | Cortex | MSA | +++ | ++ | - | - | - | +++ |
| C-0081 | Cortex | MSA | ++ | - | - | - | - | + |
| C-0030 | Cortex | MSA | ++ | + | - | + | - | ++ |
| C-0024 | Cortex | DLB | +++ | + | + | ++ | + | +++ |
| C-0031 | Cortex | DLB | ++ | + | + | + | + | ++ |
| C-0080 | Cortex | DLB | +++ | + | - | - | - | +++ |
| H-0075 | Cortex | CTRL | - | - | - | - | - | - |
| H-0294 | Cortex | CTRL | - | - | - | - | - | - |

***Antibodies used:***

**Supplementary table 1**: Antibodies used with application and dilution

| Antibody target | Supplier | Specie | Cat # | Dilution | | |
|---|---|---|---|---|---|---|
| | | | | IHC | Western blot | ELISA |
| Actin (c4) | Millipore | Mouse | MAB1501 | - | 1:10 000 | - |
| aSyn | CST | Rabbit | 2628 | 1:200 | 1:1000 | 1:5000 |
| Amytracker 520 | Ebba BioTech | - | - | 1:1000 | - | - |
| DAT | Millipore | Rat | MAB369 | 1:1000 | - | - |
| GFAP | CST | Mouse | 3670 | - | 1:1000 | - |
| Iba1 | Abcam | Rabbit | ab178846 | 1:5000 | 1:1000 | - |
| Myc tag | Abcam | Rabbit | ab9106 | 1:500 | - | - |
| Myc tag-HRP | Abcam | Mouse | ab62928 | | | 1:10 000 |
| NeuN | Millipore | Mouse | MAB377 | 1:500 | - | - |
| pS129-aSyn | Abcam | Rabbit | ab51253 | - | 1:1000 | - |
| pS129-aSyn | Custom clone 73B3 | Mouse | - | 1 ug/ml | - | - |
| pS129-aSyn | BioLegend | Mouse | 825701 | 1:1000 | - | - |
| pS129-aSyn | Wako | Mouse | 015-25191 | - | 1:2000 | 1:8000 |
| TH | Pel-Freez Biologicals | Rabbit | P40101 | 1:1000 | 1:1000 | - |
| TH | Pel-Freez Biologicals | Sheep | P60101 | 1:1000 | - | - |

***Clinical scoring system for M83 + PFF mice:***

Adapted from: Guyenet S.J., Furrer S.A., Damian V.M., Baughan T.D., La Spada A.R., Garden G.A. (2010). A simple phenotype scoring system for evaluating mouse models of cerebellar ataxia. JoVE. 39. Doi: 10.3791/1787

Each test is performed 2-3 consecutive times to ensure reproducibility. All scores are added together for a maximum clinical score of 9. Mice reaching score 8-9 were sacrificed in accordance with approved ethical protocols.

| Ledge test | | | |
|---|---|---|---|
| 0 | 1 | 2 | 3 |
| Walks on the ledge without losing its balance. Descends in the cage gracefully, using its paws. | Loses its balance and slips a couple times, able to descend in the cage without falling. | Doesn't use properly its hind legs and falls when trying to descend. | Cannot hold on to the ledge and falls. |
| Hindlimb weakness (10 sec. tail suspension) | | | |
| 0 | 1 | 2 | 3 |
| Hindlimbs consistently spayed outward, away from the abdomen. | One or both hindlimbs are not completely extended more than 50% of the time. | One hindlimb appears almost totally weak and is retracted toward the abdomen. | Both hindlimbs (and forelimbs) are completely weak, unmoving, and retracted towards the abdomen. |
| Gait | | | |
| 0 | 1 | 2 | 3 |
| Moves normally, body weight supported on all limbs, coordinated. | Weight shifting from side to side when walking, appears to limp. | Shows a severe limp, lowered pelvis and "duck feet". | Spends most of the time lying down, often on its side. |

### Supplementary figures legends

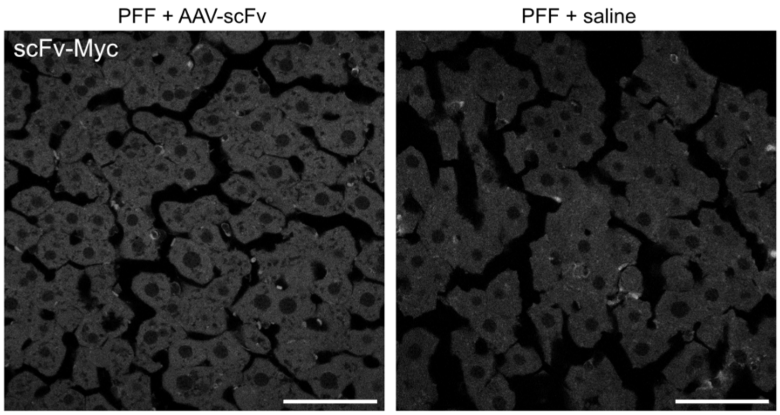


**Supplementary fig. 1: AAVCap-B10-scFv intravenous injection does not cause scFv expression in the liver at 12 weeks post-injection A)** Representative images of a Myc-tag staining in the liver of a mouse injected with AAV-scFv-73B3.

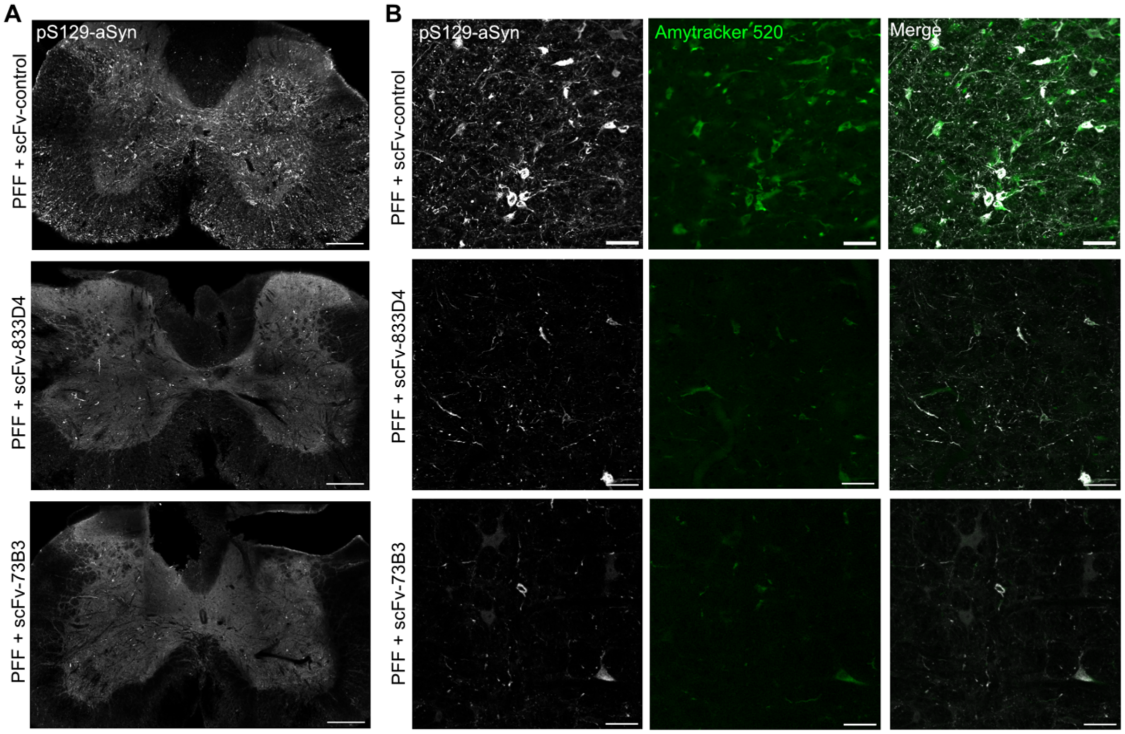


**Supplementary fig. 2: aSyn aggregates generated following PFF injection in hemizygous M83 mice are Lewy-body like and are present in the spinal cord**. **A)** Representative images showing pS129-aSyn staining in the spinal cord of PFFs-injected M83 who received scFv-control, 833D4 or 73B3. Scale bar = 200µm. **B)** Representative image showing pS129-aSyn staining colocalizing with Amytracker™ dye (β-sheets structures) 12 weeks post-PFFs injection in the medulla of an animal who received scFv-control. Scale bar = 50µm.

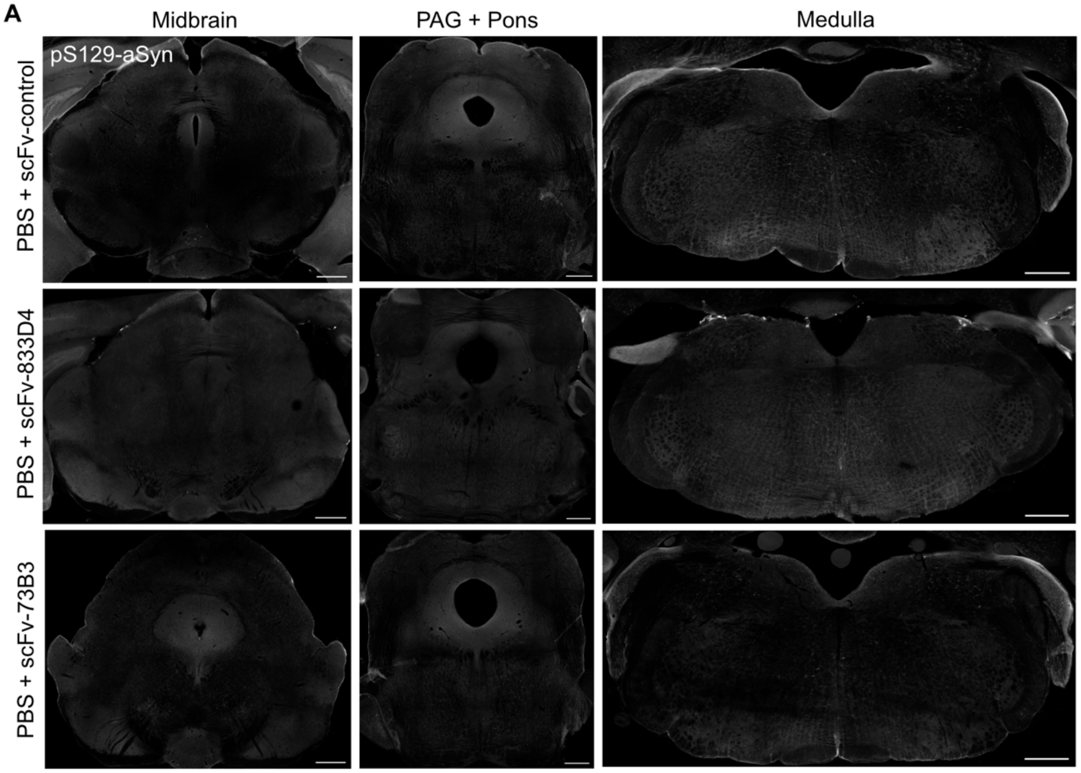


**Supplementary fig. 3: PBS injection in hemizygous M83 does not cause synucleinopathy. A)** Representative images showing pS129-aSyn staining in the midbrain, pons, PAG and medulla in mice injected with PBS and scFv control, 833D4 or 73B3. Scale bar = 500µm.

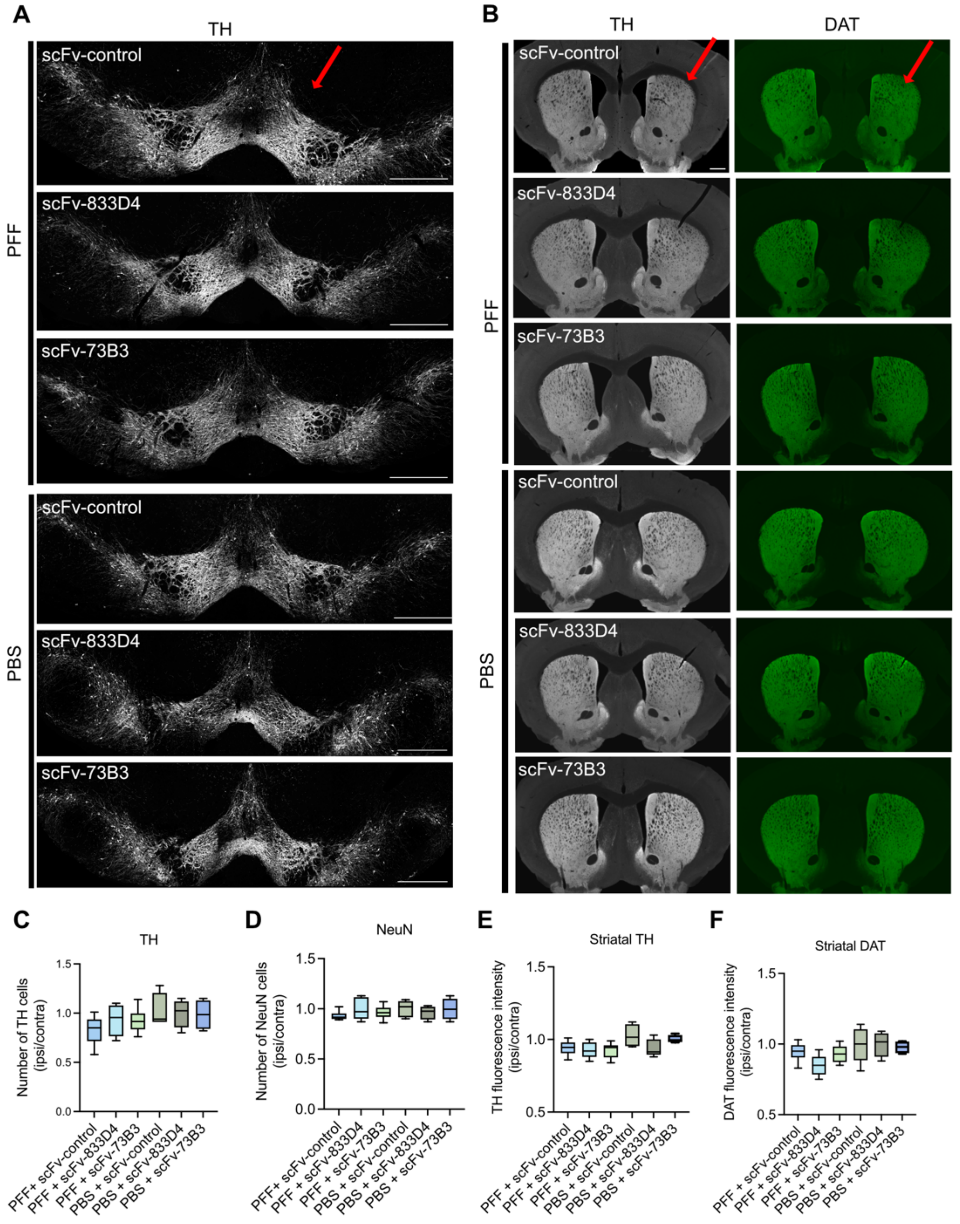


**Supplementary fig. 4: PFF injection in hemizygous M83 does not cause dopaminergic degeneration at 12 weeks post-injection. A)** Representative images of a TH immunostaining in the midbrain of a mouse from each experimental group showing no cell loss in the injected side (shown with a red arrow). **B)** Representative images of TH and DAT immunostaining in the striatum showing no axonal degeneration. **C)** Stereological counting of $TH^+$ cells in the SN and shown as a ratio of injected/uninjected side. **D)** Stereological counting of $NeuN^+$ cells in the SN and shown as a ratio of injected/uninjected side. **E)** Quantification of the fluorescence intensity of TH in the striatum and shown as a ratio of injected/uninjected side. **F)** Quantification of the fluorescence intensity of DAT in the striatum and shown as a ratio of injected/uninjected side. For C to F) n=6/group. One-way ANOVA with Dunnett's multiple comparison test.

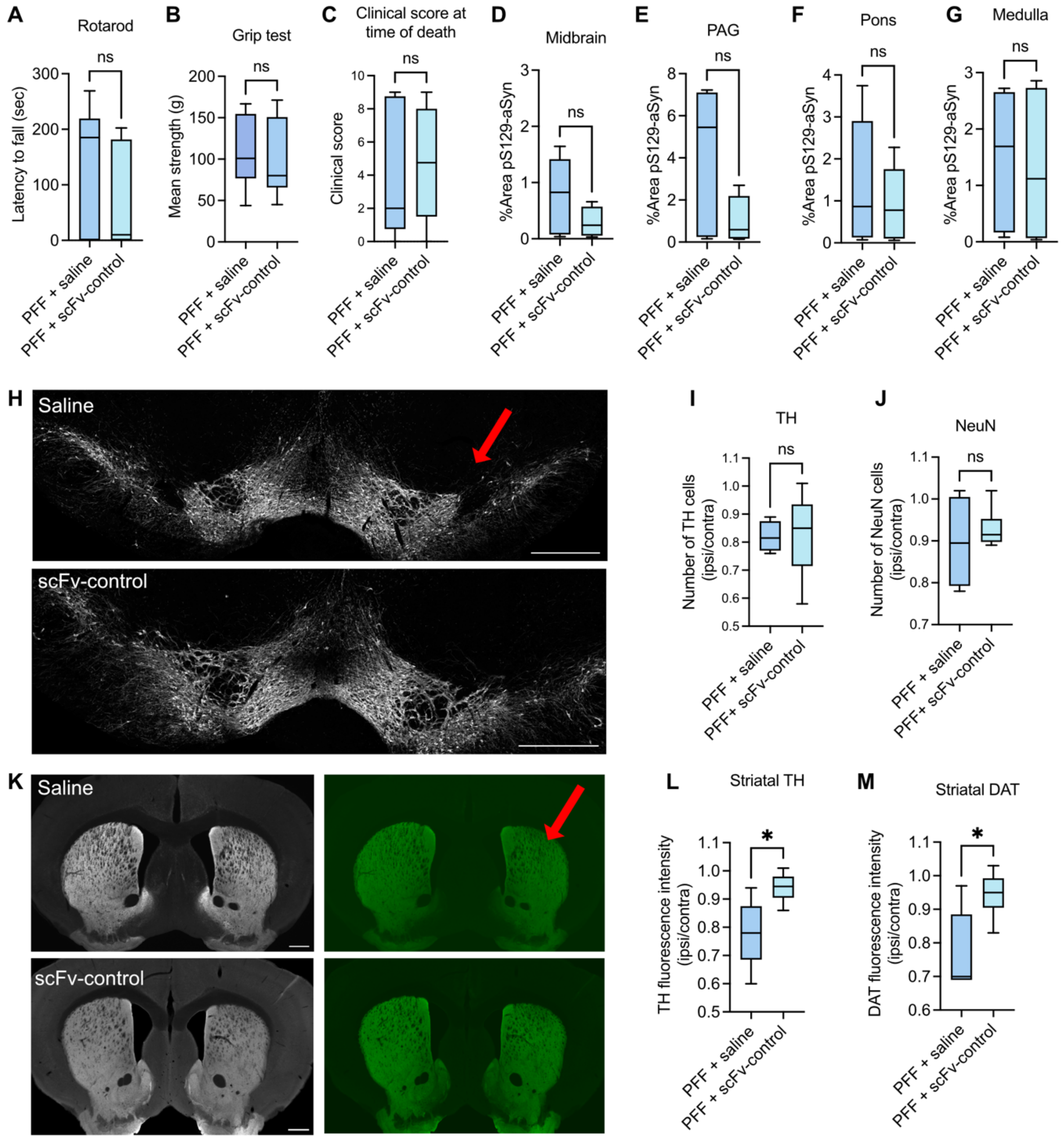


**Supplementary fig. 5: The control scFv targeting the GFP protein does not influence disease progression compared to saline injection in hemizygous M83 injected with aSyn PFF. A-C)** Comparison of mice injected with PFF and the control scFv with mice injected with saline for the rotarod (**A**), the grip test (**B**), and the clinical score at time of death **(C)** (n=9/group). Unpaired T test. **D-G)** Comparison of the 2 control groups in pS129-aSyn particles quantification in the midbrain **(D)**, the PAG **(E)**, the pons **(F)** and the medulla **(G)** (n=5-6/group). Unpaired T-test. **H)** Representative images of TH staining in the midbrain of mice injected with saline and the control scFv. **I)** Stereological count of $TH^+$ neurons in the SNc in the 2 control groups (n=5-6/group). Unpaired T-test. **J)** Stereological count of $NeuN^+$ neurons in the SNc (n=5-6/group). Unpaired T-test. **K)** Representative images of TH (white) and DAT (green) staining in the striatum. **L)** Quantification of TH intensity in the dorsal striatum (n=5-6/group). Unpaired T-test. **M)** Quantification of the DAT intensity in the dorsal striatum (n=5-6/group). Unpaired T-test $^*p<0.05$.
